# A VgrG2b fragment cleaved by caspase-11/4 promotes *Pseudomonas aeruginosa* infection through suppressing the NLRP3 inflammasome

**DOI:** 10.1101/2024.07.02.601667

**Authors:** Yan Qian, Qiannv Liu, Xiangyun Cheng, Chunlei Wang, Chun Kong, Mengqian Li, Chao Ren, Dong Jiang, Shuo Wang, Pengyan Xia

**Affiliations:** Department of Immunology, School of Basic Medical Sciences, Peking University, Beijing, 100191, China; NHC Key Laboratory of Medical Immunology, Peking University, Beijing, 100191, China; Key Laboratory of Molecular Immunology, Chinese Academy of Medical Sciences, Beijing, 100191, China; Department of Sports Medicine, Peking University Third Hospital, Beijing, 100191, China; Beijing Key Laboratory of Sports Injuries, Institute of Sports Medicine of Peking University, Beijing, 100191, China; Department of Respiratory and Critical Care Medicine, Beijing Institute of Respiratory Medicine, Beijing Chao-Yang Hospital, Capital Medical University, Beijing, 100020, China; CAS Key Laboratory of Pathogen Microbiology and Immunology, Institute of Microbiology, Chinese Academy of Sciences, Beijing, 100101, China

**Keywords:** *P. aeruginosa*, non-canonical inflammasome, caspase-11/4, VgrG2b, NLRP3

## Abstract

The T6SS of *Pseudomonas aeruginosa* plays an essential role in the establishment of chronic infections. Inflammatory cytokines mediated by inflammasomes are crucial for the body to resist bacterial infections. Here we found that during the infection of *P. aeruginosa*, non-canonical inflammasome was activated in macrophages, but the activation of downstream NLRP3 inflammasome was inhibited. The VgrG2b of *P. aeruginosa* is recognized and cleaved by caspase-11, generating a free C-terminal fragment. The VgrG2b C-terminus can bind to NLRP3, inhibiting the activation of the NLRP3 inflammasome by rejecting NEK7 binding to NLRP3. Administrating a specific peptide that inhibits the cleavage of VgrG2b by caspase-11 to mice can significantly improve their survival rate during infection. Our discovery elucidates a mechanism by which *P. aeruginosa* inhibits host immune response, providing a new approach for the future clinical treatment of *P. aeruginosa* infections.

## Introduction

Non-canonical inflammasomes are composed of caspase-11 from mice or caspase-4/5 from humans, which can recognize bacteria lipopolysaccharides (LPS) in the cytoplasm (Hagar et al., 2013; Kayagaki et al., 2013; Shi et al., 2014). Once activated, non-canonical inflammasomes induce the maturation of gasdermin D (GSDMD) to form pores on cell and mitochondrial membranes (Huang et al., 2020; Kayagaki et al., 2015; Shi et al., 2015). It also activates the NOD-, LRR-and pyrin domain-containing protein 3 (NLRP3) inflammasome (Kayagaki *et al*., 2015; Rathinam et al., 2019). NLRP3 is an important intracellular receptor that can respond to various endogenous and exogenous stimulus signals (Barnett et al., 2023; Christgen et al., 2020). NLRP3 recruits NIMA-related kinase 7 (NEK7) and undergoes polymerization, inducing the activation of caspase-1 and maturation of interleukin-1β (IL-1β) (Barnett *et al*., 2023; He et al., 2016). Non-canonical inflammasomes promote the assembly and final activation of the NLRP3 inflammasome through activating the nuclear orphan receptor NUR77 with the help of LPS and mitochondrial dsDNA (Zhu et al., 2023).

*Pseudomonas aeruginosa* is an opportunistic pathogenic bacterium widely distributed in the environment, which can cause acute and chronic infections in humans (Cendra and Torrents, 2021; Rossi et al., 2021). *P. aeruginosa* has multiple secretion systems that are used to secrete virulence factors to the external environment or host cells to promote its survival and infection (Rossi *et al*., 2021). Among them, the type III secretion system (T3SS) and the type VI secretion system (T6SS) are two important secretion systems that play important roles in acute and chronic infections of *P. aeruginosa*, respectively (Faure et al., 2014; Goodman et al., 2004). T3SS can directly inject effector proteins into target cells to alter their function or cause target cell death (Hauser, 2009; Qin et al., 2022). *P. aeruginosa* encodes three different T6SSs, namely Hcp secretion island-I (H1)-T6SS, H2-T6SS, and H3-T6SS (Hachani et al., 2011; Mougous et al., 2006). These T6SSs play a crucial role in biofilm formation, competition between bacteria, and toxicity to host cells (Chen et al., 2015; Ho et al., 2014). H1-T6SS only targets bacteria and is used for competition between bacteria (Hood et al., 2010). H2-T6SS and H3-T6SS are involved in interactions between bacteria and host cells (Jiang et al., 2014; Russell et al., 2013).

The effector proteins of H2-T6SS can target both bacteria and eukaryotic cells (Ho *et al*., 2014; Jiang *et al*., 2014). T6SS is composed of structural proteins and virulence factors (Durand et al., 2014; Ho *et al*., 2014). Structural proteins such as hemolysin-coregulated protein (Hcp), valine-glycine repeat protein G (VgrG), and proline-alanine-alanine-arginine repeats (PAAR) form a stable spike-like structure (Leiman et al., 2009; Shneider et al., 2013). Toxic factors are transported to the extracellular space through T6SS, thereby exerting toxic effects on other bacteria or host cells (Ho *et al*., 2014; Qin *et al*., 2022). VgrG2b is an important component of T6SS, which usually binds tightly to other components of T6SS such as the Hcp protein, together forming the core component of the secretion system (Sana et al., 2015). The VgrG2b protein has a special structure for recognizing and binding effector proteins, which are transported to the tip of the secretion system and injected into the target cells (Durand *et al*., 2014; Sana *et al*., 2015). VgrG2b entering eukaryotic cells can also regulate the biological functions of host cells through binding to cytoskeletal proteins (Sana *et al*., 2015; Wood et al., 2019).

Here we found that *P. aeruginosa* infection activates caspase-11 but inhibits the activation of NLRP3 inflammasomes in macrophages. The VgrG2b entering the host cell is recognized and cleaved by caspase-11 to generate a free C-terminal fragment. This fragment can bind to NLRP3 and inhibit its binding to NEK7, thereby preventing the activation of the NLRP3 inflammasome. The inhibitory effect of *P. aeruginosa* effectively avoids further activation of the host immune system and plays a crucial role in establishing bacterial infection. Our findings may provide a new perspective for the treatment of *P. aeruginosa* infections in the future.

## Results

### *P. aeruginosa* causes abnormal signaling of non-canonical inflammasome

Non-canonical inflammasomes are activated during the infection of Gram-negative bacteria such as *P. aeruginosa in vivo* (Balakrishnan et al., 2018; Kayagaki *et al*., 2013; Rathinam *et al*., 2019). Obstruction of the inflammasome pathway can lead to chronic infections (Huus et al., 2016; Phuong et al., 2021). To investigate whether T6SS of *P. aeruginosa* modulated the non-canonical inflammasome pathway to establish chronic infections, we used an *RetS* mutant strain that specifically expresses T6SS (Allsopp et al., 2017; Goodman *et al*., 2004; Han et al., 2019). As an extracellular bacterial, *P. aeruginosa* synthesizes outer membrane vesicles (OMV) to induce non-canonical inflammasome activation (Deo et al., 2020). We therefore added *P. aeruginosa* and OMVs to macrophages *in vitro* to simulate the real situation of infection *in vivo* as much as possible (Esoda and Kuehn, 2019). We found that activation of caspase-11 and cleavage of GSDMD in BMDM cells were not affected after incubation with *P. aeruginosa* and OMVs (Fig. 1A, B). However, activation of caspase-1 was significantly inhibited in cells infected with *P. aeruginosa* (Fig. 1A). Consistent with the results of immunoblotting, the secretion of IL-1β was diminished after infection with *P. aeruginosa* (Fig. 1C). Although the mortality rate of *P. aeruginosa*-infected cells was increased (Fig. 1D) and the survival rate was decreased (Fig. 1E), there was not much difference compared to cells stimulated with intracellular *E. coli*. These results indicate that the activation of NLRP3 inflammasomes post non-canonical inflammasome stimulation is inhibited in the presence of *P. aeruginosa*. Similar results were observed in human macrophage experiments, with normal caspase-4 activation and GSDMD cleavage after *P. aeruginosa* infection (Fig. S1A, B). However, the activation of caspase-1 was suppressed and IL-1β could not be secreted normally in these cells (Fig. S1A, C). With *P. aeruginosa* infection, the cell mortality rate was increased (Fig. S1D) and the survival rate was decreased (Fig. S1E), showing no much difference with cells treated only with intracellular *E. coli*.

**Figure 1.**
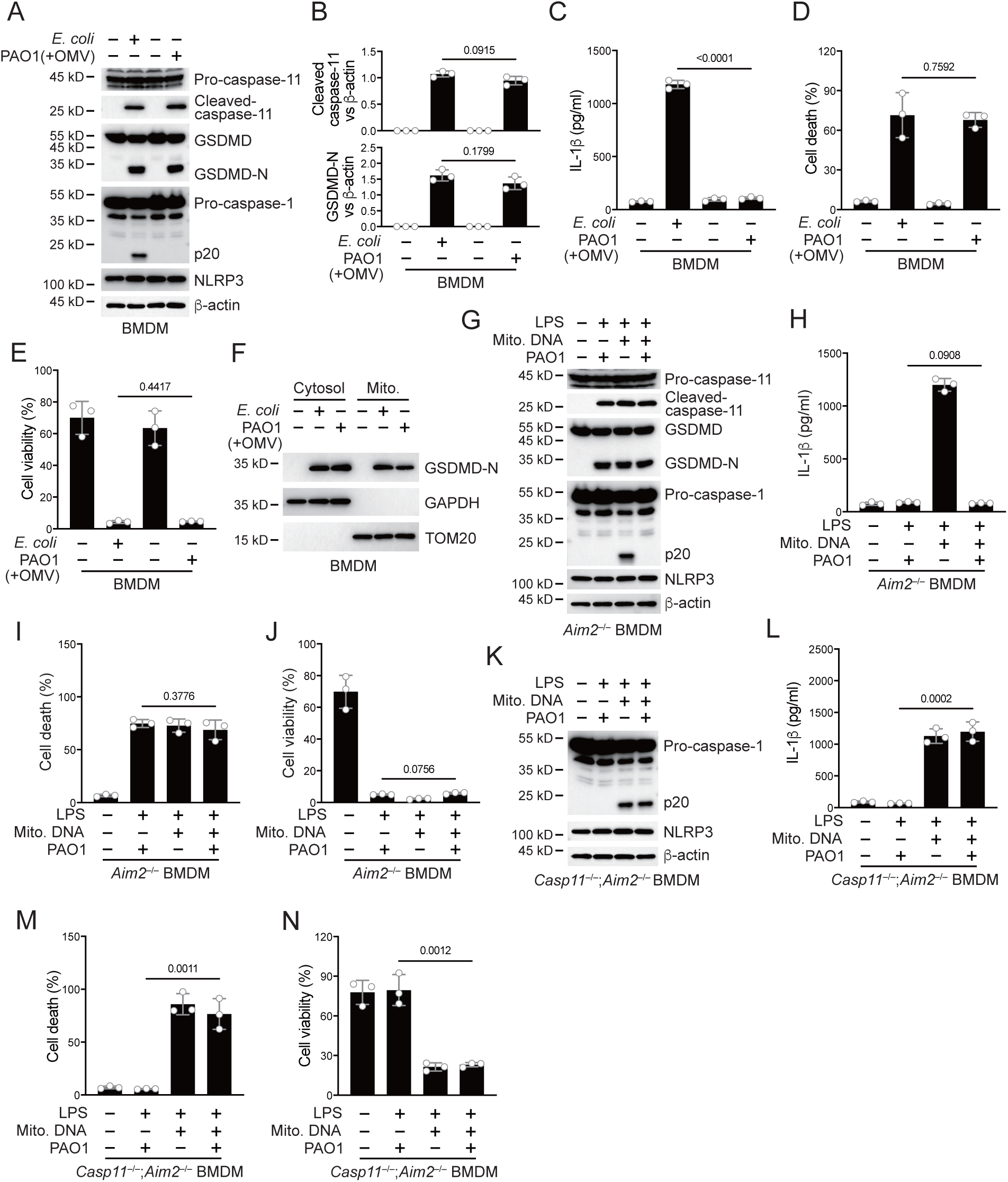
*P. aeruginosa* infection suppresses NLRP3 inflammasome. (A–E) Wild-type BMDM cells were primed overnight with 1 μg/ml Pam3CSK4 and incubated with *E. coli* or *P. aeruginosa* (ΔRetS PAO1) at an MOI of 30 and 20 μg/ml OMVs for 2 h. Cells were then supplemented with fresh medium containing 100 μg/ml Gentamycin. Cells were lysed and immunoblotted as indicated 16 h post infection (A). Band intensities of cleaved caspase-11 (top) and GSDMD (bottom) were quantified and compared to that of β-actin (B). Cell culture supernatants were collected for an ELISA assay to determine the secreted IL-1β protein levels 16 h post infection (C). Cytotoxicity was determined by LDH release assay in cell culture supernatants 16 h post infection (D). Cell viability was determined by an ATP quantification assay in cell pellets 16 h post infection (E). (F) Wild-type BMDM cells were primed overnight with 1 μg/ml Pam3CSK4 and incubated with *E. coli* or ΔRetS PAO1 at an MOI of 30 and 20 μg/ml OMVs for 2 h. Cells were then supplemented with fresh medium containing 100 μg/ml Gentamycin. Cells were subjected to cellular component fractionation and immunoblotted as indicated 16 h post infection. (G–J) *Aim2*^−/−^ BMDM cells were primed overnight with 1 μg/ml Pam3CSK4, followed by transfection of 2 μg/ml LPS and 2 μg/ml extracted BMDM mitochondrial DNA using DOTAP with or without the incubation of ΔRetS PAO1 at an MOI of 30 for 2 h. Cells were then supplemented with fresh medium containing 100 μg/ml Gentamycin. Cells were lysed and immunoblotted 16 h post infection (G). Cell culture supernatants were collected for an ELISA assay to determine the secreted IL-1β protein levels 16 h post infection (H). Cytotoxicity was determined by LDH release assay in cell culture supernatants 16 h post infection (I). Cell viability was determined by an ATP quantification assay in cell pellets 16 h post infection (J). (K–N) *Casp11*^−/−^;*Aim2*^−/−^ BMDM cells were primed overnight with 1 μg/ml Pam3CSK4, followed by transfection of 2 μg/ml LPS and 2 μg/ml extracted BMDM mitochondrial DNA using DOTAP with or without the incubation of ΔRetS PAO1 at an MOI of 30 for 2 h. Cells were then supplemented with fresh medium containing 100 μg/ml Gentamycin. Cells were lysed and immunoblotted 16 h post infection (K). Cell culture supernatants were collected for an ELISA assay to determine the secreted IL-1β protein levels 16 h post infection (L). Cytotoxicity was determined by LDH release assay in cell culture supernatants 16 h post infection (M). Cell viability was determined by an ATP quantification assay in cell pellets 16 h post infection (N). Data were shown as means±SD. For1C, 1D, 1E, 1H, 1I, 1J, 1L, 1M and 1N, data of three independent experiments were calculated. Experiments were repeated three times with similar results.

There might be two mechanisms by which NLRP3 inflammasomes were inhibited during this process. One was that the NLRP3 inflammasome lacked necessary stimuli for activation, and the other was that the NLRP3 inflammasome was inhibited by a bacterial substance. After the activation of caspase-11, GSDMD pores can be detected in mitochondrial membranes, causing the release of mtDNA and ROS which amplify NLRP3 activation (Miao et al., 2023; Xian et al., 2022). We first detected the localization of GSDMD N-terminus on the mitochondrial membrane during *P. aeruginosa* infection. The results indicate that GSDMD-N can be normally localized on the mitochondrial membrane (Fig. 1F). In addition, the release of mitochondrial DNA into the cytoplasm was not affected by *P. aeruginosa* infection (Fig. S1F). The binding of NUR77 and NLRP3 was not affected during the infection of *P. aeruginosa* (Fig. S1G). However, the binding of NLRP3 to NEK7 was significantly inhibited (Fig. S1H). These data suggest that the inhibition of NLRP3 is potentially influenced by a certain inhibitory factor from *P. aeruginosa*.

To verify this, we introduced LPS and mitochondrial DNA into macrophages to induce NLRP3 inflammasome activation, bypassing the caspase-11-GSDMD prerequisite (Zhu *et al*., 2023). To avoid the activation of the AIM2 inflammasome by mitochondrial DNA, we conducted the following experiments under an *Aim2* knockout background. We found that after introducing LPS and mitochondrial DNA, NLRP3 inflammasomes were activated (Fig. 1G). However, NLRP3 activation was inhibited (Fig. 1G) and the secretion of IL-1β was tremendously inhibited (Fig. 1H) after incubation with *P. aeruginosa*. The percentage of cells undergoing pyroptosis (Fig. 1I) and the cell viability (Fig. 1J) did not change due to infection with *P. aeruginosa*. This was consistent with previous reports suggesting that GSDMD cleavage by caspase-11, rather than subsequent GSDMD cleavage by NLRP3, is the main factor causing cell death (Broz et al., 2020; Kayagaki et al., 2011). Next, we used *Gsdmd* and *Aim2* double knockout cells to detect the effect of *P. aeruginosa* on NLRP3 inflammasome activation. We found that after *Gsdmd* knockout, the NLRP3 inflammasome became active when LPS and mitochondrial DNA were introduced into macrophages. However, NLRP3 was still inhibited after incubation with *P. aeruginosa* (Fig. S1I). The secretion of IL-1β was also inhibited due to *Gsdmd* deficiency (Fig. S1J). The cell death rate was low due to the absence of GSDMD (Fig. S1K), while the percentage of viable cells was relatively high (Fig. S1L). These results suggest that potent inhibitory factors of *P. aeruginosa* may act upstream of GSDMD. Therefore, we conducted infection experiments using *Casp11* and *Aim2* double knockout cells. After incubation with *P. aeruginosa*, the activation of NLRP3 inflammasome was not affected (Fig. 1K). The secretion of IL-1β was no longer inhibited (Fig. 1L). And the cell death rate was high due to the processing of GSDMD by activated caspase-1 (Fig. 1M), while the survival rate was low (Fig. 1N). These data indicate that the inhibitory factors of *P. aeruginosa* requires caspase-11 to take effect.

### Caspase-11 cleaves VgrG2b of *P. aeruginosa*

In order to search for potential substrates of caspase-11 in *P. aeruginosa*, we designed a systematic screening system (Fig. 2A). Simply, we constructed two fluorescent plasmids, one containing green fluorescent protein (GFP), polyclonal sites, and mitochondrial outer membrane localization signals, and the other containing red fluorescent protein (RFP) and mitochondrial outer membrane localization signals. We extracted the mRNA of *P. aeruginosa* and prepared it into a library for insertion into GFP containing plasmids. We co-transfected it with the RFP plasmid into HEK293T cells and sorted the monoclonal cells containing both RFP and GFP signals into a 96-well plate using flow cytometry. Subsequently, the plasmid encoding caspase-11 was transfected into cells, and cells with disappearance of green signal on mitochondria were selected. The target proteins in these cells were cleaved by caspase-11, causing the detachment of GFP from mitochondria. After sequencing, we found that VgrG1a and VgrG2b repeatedly appeared in cells that met the screening criteria.

**Figure 2.**
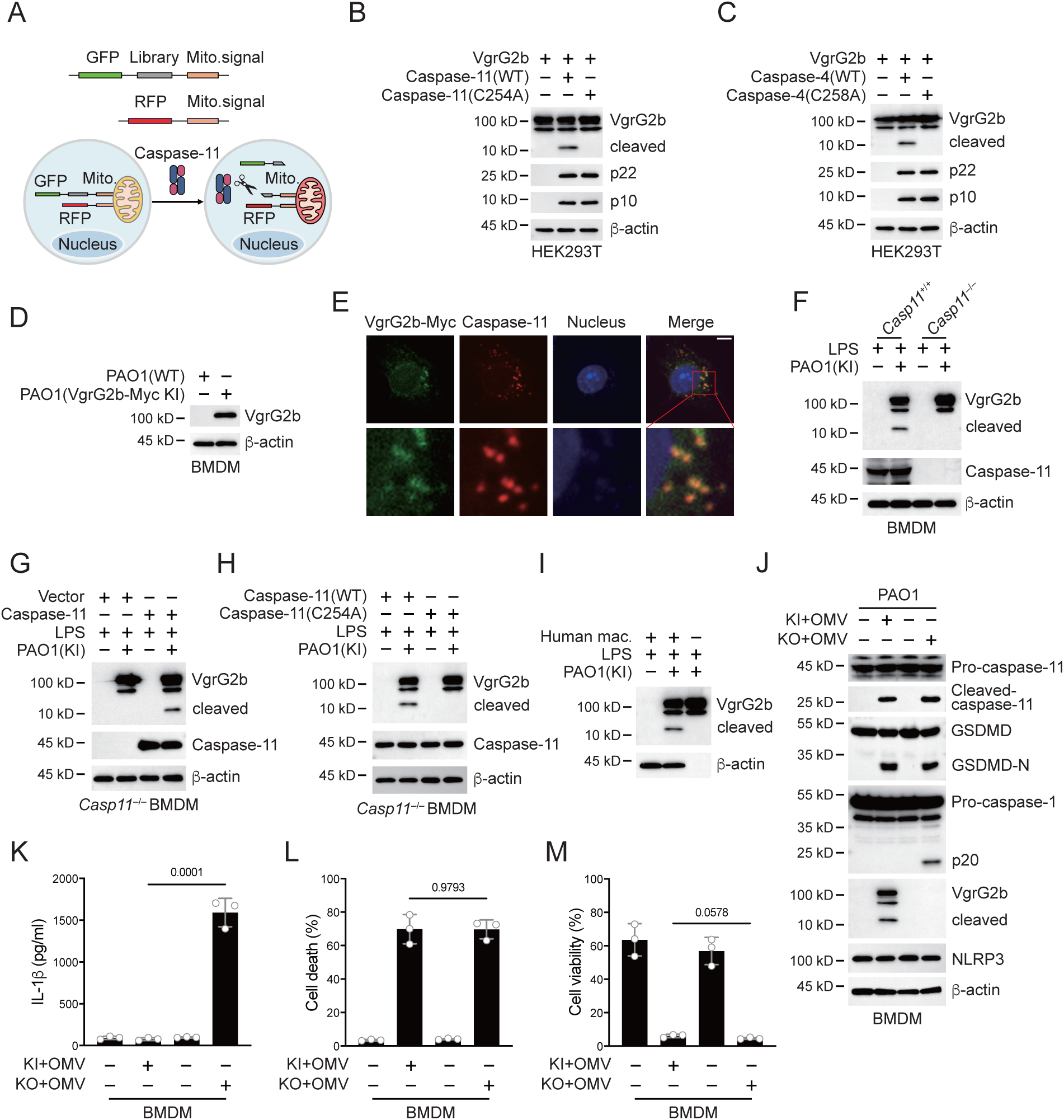
*P. aeruginosa* VgrG2b is cleaved by caspase-11. (A) Screening strategy for identifying caspase-11 substrates in *P. aeruginosa*. Briefly, *P. aeruginosa* cDNAs were cloned into GFP vectors containing a mitochondrial localization signal. RFP vectors with a mitochondrial localization signal were used as transfection controls. HEK293T cells were co-transfected with these vectors and caspase-11. Cells losing GFP signals on mitochondria were sequenced. (B, C) Plasmids encoding VgrG2b and mouse caspase-11 p22/p10 (B) or human caspase-4 p22/p10 (C) were co-transfected into HEK293T cells for 24 h, followed by immunoblotting with antibodies against the indicated proteins. (D) Wild-type BMDM cells were primed overnight with 1 μg/ml Pam3CSK4 and incubated with ΔRetS PAO1 or VgrG2b-Myc knockin ΔRetS PAO1 at an MOI of 30 and 20 μg/ml OMVs for 2 h. Cells were then supplemented with fresh medium containing 100 μg/ml Gentamycin. Cells were lysed and immunoblotted as indicated 16 h post infection. (E) Wild-type BMDM cells were primed overnight with 1 μg/ml Pam3CSK4 and incubated with VgrG2b-Myc knockin ΔRetS PAO1 at an MOI of 30 and 20 μg/ml OMVs for 2 h. Cells were then supplemented with fresh medium containing 100 μg/ml Gentamycin. Cells were fixed and stained with antibodies against Myc or caspase-11 16 h post infection. Nucleus was stained with DAPI. Scale bar, 5 μm. (F) *Casp11*^+/+^ and *Casp11*^−/−^ BMDM cells were primed overnight with 1 μg/ml Pam3CSK4, followed by transfection of 2 μg/ml LPS using DOTAP with or without the incubation of VgrG2b-Myc knockin ΔRetS PAO1 at an MOI of 30 for 2 h. Cells were then supplemented with fresh medium containing 100 μg/ml Gentamycin. Cells were lysed and immunoblotted 16 h post infection. (G) *Casp11*^−/−^ BMDM cells were infected with lentiviruses encoding a control vector of caspase-11. Cells were primed overnight with 1 μg/ml Pam3CSK4, followed by transfection of 2 μg/ml LPS using DOTAP with or without the incubation of VgrG2b-Myc knockin ΔRetS PAO1 at an MOI of 30 for 2 h. Cells were then supplemented with fresh medium containing 100 μg/ml Gentamycin. Cells were lysed and immunoblotted as indicated 16 h post infection. (H) *Casp11*^−/−^ BMDM cells were infected with lentiviruses encoding wild-type or mutant caspase-11. Cells were primed overnight with 1 μg/ml Pam3CSK4, followed by transfection of 2 μg/ml LPS using DOTAP with or without the incubation of VgrG2b-Myc knockin ΔRetS PAO1 at an MOI of 30 for 2 h. Cells were then supplemented with fresh medium containing 100 μg/ml Gentamycin. Cells were lysed and immunoblotted as indicated 16 h post infection. (I) Human macrophages were primed overnight with 1 μg/ml Pam3CSK4, followed by transfection of 2 μg/ml LPS using DOTAP with or without the incubation of VgrG2b-Myc knockin ΔRetS PAO1 at an MOI of 30 for 2 h. Cells were then supplemented with fresh medium containing 100 μg/ml Gentamycin. Cells were lysed and immunoblotted as indicated 16 h post infection. (J– M) Wild-type BMDM cells were primed overnight with 1 μg/ml Pam3CSK4, followed by incubation of VgrG2b-Myc knockin (KI) or knockout (KO) ΔRetS PAO1 at an MOI of 30 and 20 μg/ml OMVs for 2 h. Cells were then supplemented with fresh medium containing 100 μg/ml Gentamycin. Cells were lysed and immunoblotted as indicated 16 h post infection (J). Cell culture supernatants were collected for an ELISA assay to determine the secreted IL-1β protein levels 16 h post infection (K). Cytotoxicity was determined by LDH release assay in cell culture supernatants 16 h post infection (L). Cell viability was determined by an ATP quantification assay in cell pellets 16 h post infection (M). Data were shown as means±SD. For 2K, 2L and 2M, data of three independent experiments were calculated. Experiments were repeated three times with similar results.

We cloned the genes of VgrG1a and VgrG2b and cleaved them using mouse caspase-11. We found that they can both be cleaved (Fig. S2A and Fig. 2B). We also found that they can be cleaved by human caspase-4 (Fig. S2B and Fig. 2C), but not by caspase-11/4 mutants lacking enzymatic activities. These results indicate that our screening system is reliable. The number of target genes screened was very limited, which was consistent with the particularly narrow substrate spectrum of caspae-11 (Shi et al., 2023). We also tested other members of the VgrG family and found that neither mouse caspase-11 nor human caspase-4 could cleave these members (Fig. S2C–L). VgrG1a and VgrG2b belong to the secretion systems of H1-T6SS and H2-T6SS, and H1-T6SS does not have an effect on eukaryotic cells (Hood *et al*., 2010; Sana *et al*., 2015). Therefore, our subsequent research focused on VgrG2b.

Our next concern was whether VgrG2b entered macrophages. We found that after co-culturing of *P. aeruginosa* with BMDM cells, the VgrG2b protein appeared in macrophages (Fig. 2D). Immunofluorescence experiments also showed the presence of bacterial proteins in macrophages after co-incubation with *P. aeruginosa* (Fig. 2E). We then incubated *P. aeruginosa* with wild-type BMDM cells and found that VgrG2b was cleaved in macrophages, while the cleavage band of VgrG2b disappeared in *Casp11*^−/−^ BMDM cells (Fig. 2F). We restored caspase-11 expression in *Casp11*^−/−^ BMDM cells and found that the presence of caspase-11 induced VgrG2b cleavage (Fig. 2G). We also rescued *Casp11*^−/−^ BMDM cells with wild-type and enzymatic mutant caspase-11 and found that mutant caspase-11 could not induce VgrG2b cleavage (Fig. 2H). In addition, we incubated human macrophages with *P. aeruginosa*. As expected, VgrG2b was cleaved in human cells (Fig. 2I). Finally, to verify that *P. aeruginosa* affected inflammasome activation through H2-T6SS, we generated a *P. aeruginosa* strain with VgrG2b deficiency. We found that *P. aeruginosa* lacking VgrG2b no longer inhibited the NLRP3 inflammasome (Fig. 2J and Fig. S2M) and IL-1β could be secreted normally in the mutant bacteria-treated cells (Fig. 2K). However, the percentage of cell death was not affected (Fig. 2L) and the level of cell viability remained unchanged among mutants and wild-type controls (Fig. 2M). These data indicate that H2-T6SS plays an important role in inhibiting host immune response through VgrG2b during *P. aeruginosa* infection, primarily targeting NLRP3 inflammasome activation rather than caspase-11 activation and GSDMD cleavage.

### VgrG2b cleavage at its C-terminus is crucial for NLRP3 inhibition

Caspase-11 recognizes its substrate through a tertiary structure formed by its p22/p10 fragments (Wang et al., 2020). To further verify that VgrG2b was a substrate of caspase-11, we co-expressed enzymatically inactive mutant caspase-11 p22/p10 and VgrG2b in HEK293T cells. Co-immunoprecipitation results showed that VgrG2b bound to the p22/p10 complex (Fig. 3A). VgrG2b also bound to the p22/p10 complex of mutant human caspase-4 (Fig. S3A). These results suggest that VgrG2b can indeed be recognized by caspase-11/4. Based on the size of the cleaved VgrG2b fragment, we speculated that the cleavage site should be at its C-terminus. We constructed mutant plasmids carrying aspartic acid substitutions for VgrG2b and found that only the D883A mutant was no longer cleaved by caspase-11 (Fig. 3B). We also verified that this VgrG2b mutant could not be cleaved by human caspase-4 (Fig. 3C). Meanwhile, we constructed a chimeric protein containing VgrG2a N-terminus, a linker sequence around D883 and VgrG2b C-terminus (Fig. S3B). This chimeric protein could be cleaved by caspase-11 (Fig. 3D). Another chimeric protein containing VgrG2b N-terminus, a linker sequence around D883 and GFP cannot be cleaved by caspase-11 (Fig. S3B, C). Based on the substrate cleavage characteristic of caspase-11 that it needs to bind the target first to mediate cleavage (Wang *et al*., 2020), we speculated that the VgrG2b C-terminus was the site where VgrG2b bound to caspase-11. Therefore, we tested the binding of VgrG2b C-terminus with caspase-11. Indeed, the C-terminus of VgrG2b bound to the p22/p10 complex (Fig. 3E). Similarly, VgrG2b C-terminus also bound to p22/p10 of human caspase-4 (Fig. S3D). VgrG2b was recombinantly expressed and incubated with the p22/p10 complex. We found that caspase-11 induced the cleavage of the recombinant VgrG2b (Fig. 3F and Fig. S3E).

**Figure 3.**
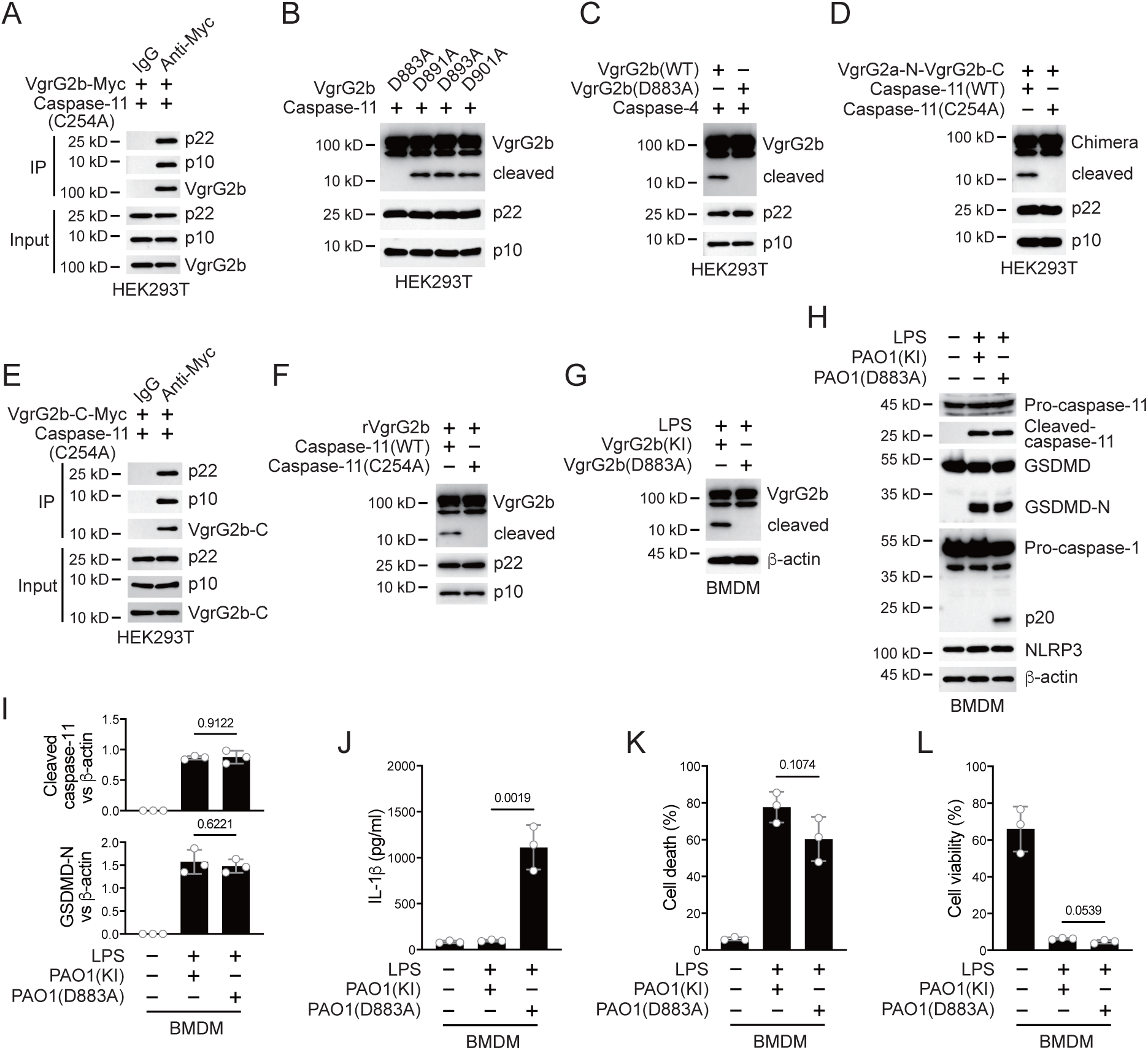
Caspase-11 cleaves VgrG2b at D883. (A) Plasmids encoding VgrG2b and mutant caspase-11 p22/p10 were co-transfected into HEK293T cells for 24 h, followed by immunoprecipitation with a control IgG or antibody against Myc. Precipitates were immunoblotted as indicated. (B) Plasmids encoding caspase-11 p22/p10 and VgrG2b mutants were co-transfected into HEK293T cells for 24 h, followed by immunoblotting with antibodies against the indicated proteins. (C) Plasmids encoding caspase-4 p22/p10 and VgrG2b variants were co-transfected into HEK293T cells for 24 h, followed by immunoblotting with antibodies against the indicated proteins. (D) Plasmids encoding VgrG2a-N-VgrG2b-C chimera and caspase-11 p22/p10 were co-transfected into HEK293T cells for 24 h, followed by immunoblotting with antibodies against the indicated proteins. (E) Plasmids encoding VgrG2b C-terminus and mutant caspase-11 p22/p10 were co-transfected into HEK293T cells for 24 h, followed by immunoprecipitation with a control IgG or antibody against Myc. Precipitates were immunoblotted as indicated. (F) Recombinant VgrG2b were incubated with caspase-11 p22/p10 for 4 h, followed by immunoblotting with antibodies against the indicated proteins. (G) Wild-type BMDM cells were primed overnight with 1 μg/ml Pam3CSK4, followed by transfection of 2 μg/ml LPS using DOTAP with or without the incubation of VgrG2b wild-type (KI) or mutant (D883A) knockin ΔRetS PAO1 at an MOI of 30 for 2 h. Cells were then supplemented with fresh medium containing 100 μg/ml Gentamycin. Cells were lysed and immunoblotted 16 h post infection. (H–L) Wild-type BMDM cells were primed overnight with 1 μg/ml Pam3CSK4, followed by transfection of 2 μg/ml LPS using DOTAP with or without the incubation of VgrG2b wild-type (KI) or mutant (D883A) knockin ΔRetS PAO1 at an MOI of 30 for 2 h. Cells were then supplemented with fresh medium containing 100 μg/ml Gentamycin. Cells were lysed and immunoblotted as indicated 16 h post infection (H). Band intensities of cleaved caspase-11 (top) and GSDMD (bottom) were quantified and compared to that of β-actin (I). Cell culture supernatants were collected for an ELISA assay to determine the secreted IL-1β protein levels 16 h post infection (J). Cytotoxicity was determined by LDH release assay in cell culture supernatants 16 h post infection (K). Cell viability was determined by an ATP quantification assay in cell pellets 16 h post infection (L). Data were shown as means±SD. For 3J, 3K and 3L, data of three independent experiments were calculated. Experiments were repeated three times with similar results.

We generated a *P. aeruginosa* strain with a D883A mutation in the VgrG2b gene. After incubating this bacterium with mouse BMDM, mutant VgrG2b did not undergo cleavage (Fig. 3G). Consistently, the NLRP3 inflammasome was no longer inhibited (Fig. 3H, I) and the secretion of IL-1β was restored post incubation with this *P. aeruginosa* mutant (Fig. 3J), indicating that VgrG2b cleavage by caspase-11 is crucial for NLRP3 inhibition. The cell death and survival rates were not affected (Fig. 3K, L). Moreover, inhibition of caspase-11 but not caspase-3 or caspase-7 blocked VgrG2b cleavage (Fig. S3F). Similarly, after incubation of the *P. aeruginosa* mutant with human macrophages, the NLRP3 inflammasome was not suppressed (Fig. S3G, H) and IL-1β was secreted normally (Fig. S3I). The cell death and survival rates were not affected (Fig. S3J, K). These results indicate that VgrG2b cleavage at its C-terminus is crucial for NLRP3 inhibition.

### VgrG2b C-terminus interacts with NLRP3

In order to investigate how VgrG2b C-terminus inhibited the NLRP3 inflammasome, we performed a yeast two-hybrid screening using the C-terminus of VgrG2b as a bait and screened a mouse bone marrow cDNA library. In the positive clones obtained from the screening, the NLRP3 fragments repeatedly appeared (Fig. 4A). We validated their interaction in HEK293T cells and found that VgrG2b C-terminus indeed bound to NLRP3 (Fig. 4B). Full-length VgrG2b also bound to NLRP3 (Fig. 4C), but the N-terminus of VgrG2b did not bind to NLRP3 (Fig. S4A). VgrG2b C-terminus was introduced into BMDM cells through cell-penetrating peptides, and it was found that the C-terminus interacted with endogenous NLRP3 (Fig. 4D). Similarly, VgrG2b C-terminus also interacted with the human NLRP3 (Fig. S4B). In addition, we incubated *P. aeruginosa* with mouse BMDM cells and found that NLRP3 interacted with the C-terminus of VgrG2b rather than the full-length one (Fig. 4E). The same result was observed after incubation of *P. aeruginosa* with human macrophages, where NLRP3 bound to the C-terminus of VgrG2b (Fig. S4C). The colocalization between VgrG2b C-terminus and NLRP3 was observed inside BMDM cells (Fig. 4F). The VgrG2b C-terminus did not bind to mouse GSDMD (Fig. S4D), ASC (Fig. S4E), nor caspase-1 (Fig. S4F, G), indicating that the binding targets of VgrG2b in host cells is very specific. We further searched for the region where VgrG2b C-terminus interacted with NLRP3 (Fig. 4G), and found that their interaction area was at the NACHT-LRR junction of NLRP3 (Fig. 4H, I).

**Figure 4.**
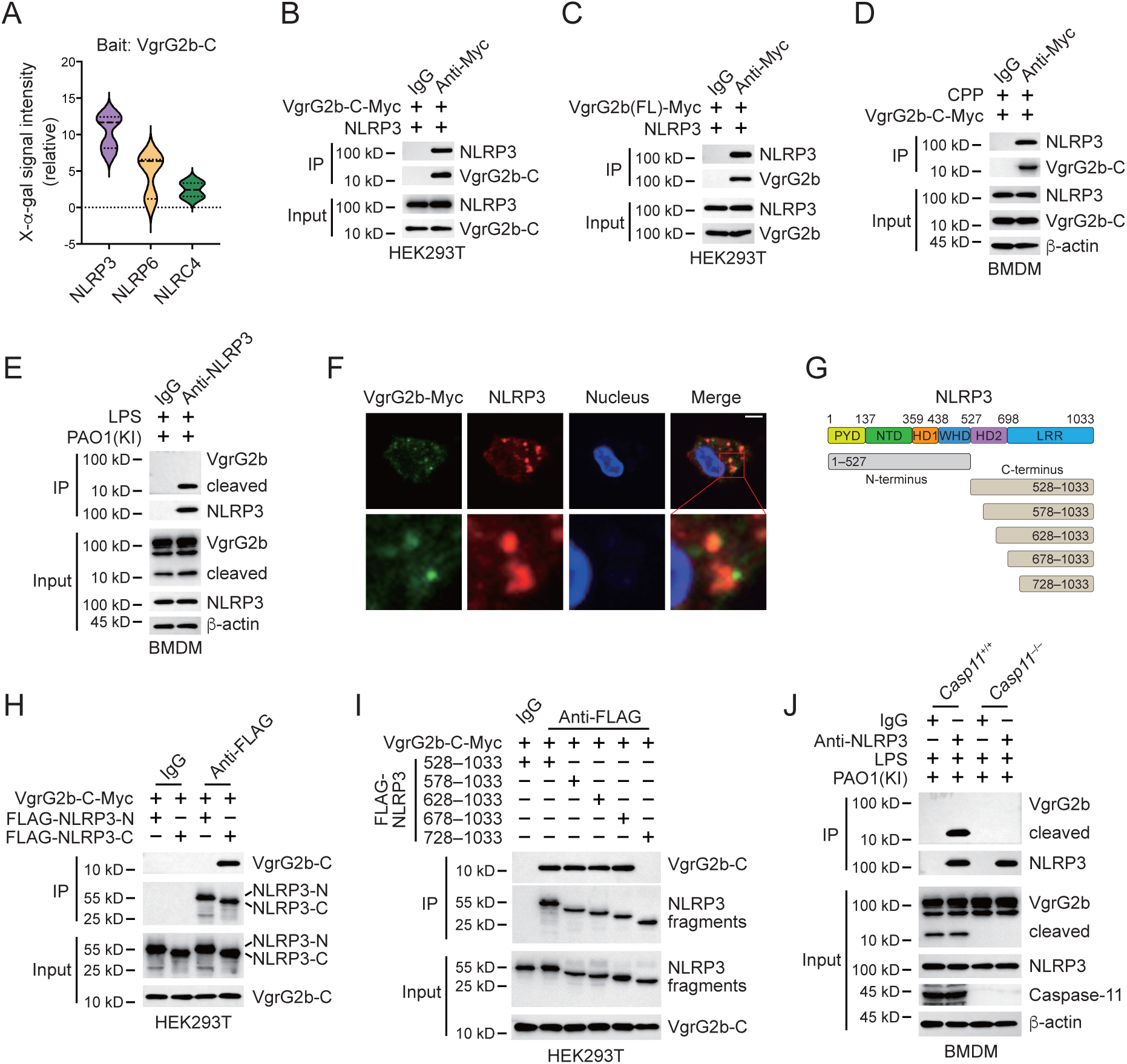
VgrG2b C-terminus interacts with NLRP3. (A) Yeast two-hybrid screening was performed using VgrG2b C-terminus as bait and a mouse bone marrow cDNA library was screened. The interaction strength between bait and preys were visualized by X-α-gal assays. (B, C) Plasmids encoding VgrG2b C-terminus (B) or full-length (C) and NLRP3 were co-transfected into HEK293T cells for 24 h, followed by immunoprecipitation with a control IgG or antibody against Myc. Precipitates were immunoblotted as indicated. (D) VgrG2b C-terminus proteins were introduced into BMDM cells with the help of cell-penetrating peptides (CPP) for 6 h. Cells were lysed and immunoprecipitated with a control IgG or antibody against Myc. Precipitates were immunoblotted as indicated. (E) BMDM cells were primed overnight with 1 μg/ml Pam3CSK4, followed by transfection of 2 μg/ml LPS using DOTAP with the incubation of VgrG2b knockin (KI) ΔRetS PAO1 at an MOI of 30 for 2 h. Cells were then supplemented with fresh medium containing 100 μg/ml Gentamycin. Cells were lysed and immunoprecipitated with a control IgG or antibody against NLRP3 16 h post infection. Precipitates were immunoblotted as indicated. (F) BMDM cells were primed overnight with 1 μg/ml Pam3CSK4, followed by transfection of 2 μg/ml LPS using DOTAP with the incubation of VgrG2b knockin (KI) ΔRetS PAO1 at an MOI of 30 for 2 h. Cells were then supplemented with fresh medium containing 100 μg/ml Gentamycin. Cells were fixed and stained with antibodies against Myc and NLRP3 16 h post infection. Nucleus was stained with DAPI. Scale bar, 5 μm. (G) Scheme for NLRP3 truncations. (H, I) Plasmids encoding VgrG2b C-terminus and NLRP3 truncations were co-transfected into HEK293T cells for 24 h, followed by immunoprecipitation with a control IgG or antibody against FLAG. Precipitates were immunoblotted as indicated. (J) *Casp11*^+/+^ and *Casp11*^−/−^ BMDM cells were primed overnight with 1 μg/ml Pam3CSK4, followed by transfection of 2 μg/ml LPS using DOTAP with the incubation of VgrG2b-Myc knockin (KI) ΔRetS PAO1 at an MOI of 30 for 2 h. Cells were then supplemented with fresh medium containing 100 μg/ml Gentamycin. Cells were lysed and immunoprecipitated with a control IgG or antibody against NLRP3 16 h post infection. Precipitates were immunoblotted as indicated. Data were shown as means±SD. Experiments were repeated three times with similar results.

In *Casp11* knockout BMDM cells, VgrG2b could not be cleaved and did not bind to NLRP3 (Fig. 4J). However, in *Gsdmd* knockout cells, VgrG2b still underwent cleavage and bound to NLRP3 (Fig. S4H). In the absence of GSDMD, the NLRP3 inflammasome was not activated (Broz *et al*., 2020; Kayagaki *et al*., 2015). This result indicates that the binding of VgrG2b C-terminus to NLRP3 is independent of NLRP3 activation, but requires a cleaved segment of VgrG2b processed by caspase-11. VgrG2b also possesses a metalloprotease activity (Wood *et al*., 2019). To test whether its protease activity is responsible for VgrG2b cleavage observed in our study, we mutated VgrG2b to disable its protease activity and found that this mutant form did not prohibit caspase-11 cleavage on VgrG2b (Fig. S4I). Moreover, VgrG2b mutant C-terminus bound NLRP3 and inhibited NLRP3 activation just like its wild-type counterpart (Fig. S4J, K).

In all, these results suggest that the cleaved VgrG2b C-terminus binds NLRP3.

### VgrG2b C-terminus suppresses the NLRP3 inflammasome

Whether VgrG2b C-terminus directly inhibited the activation of NLRP3 inflammasomes was the focus of our next exploration. We validated this hypothesis in the HEK293T recombination system. The VgrG2b C-terminus significantly inhibited the activation of NLRP3 inflammasome originated from mouse (Fig. 5A) or human (Fig. S5A). In addition, overexpression of VgrG2b C-terminus suppressed the activation of classical NLRP3 inflammasomes in both mouse (Fig. 5B) and human macrophages (Fig. S5B).

**Figure 5.**
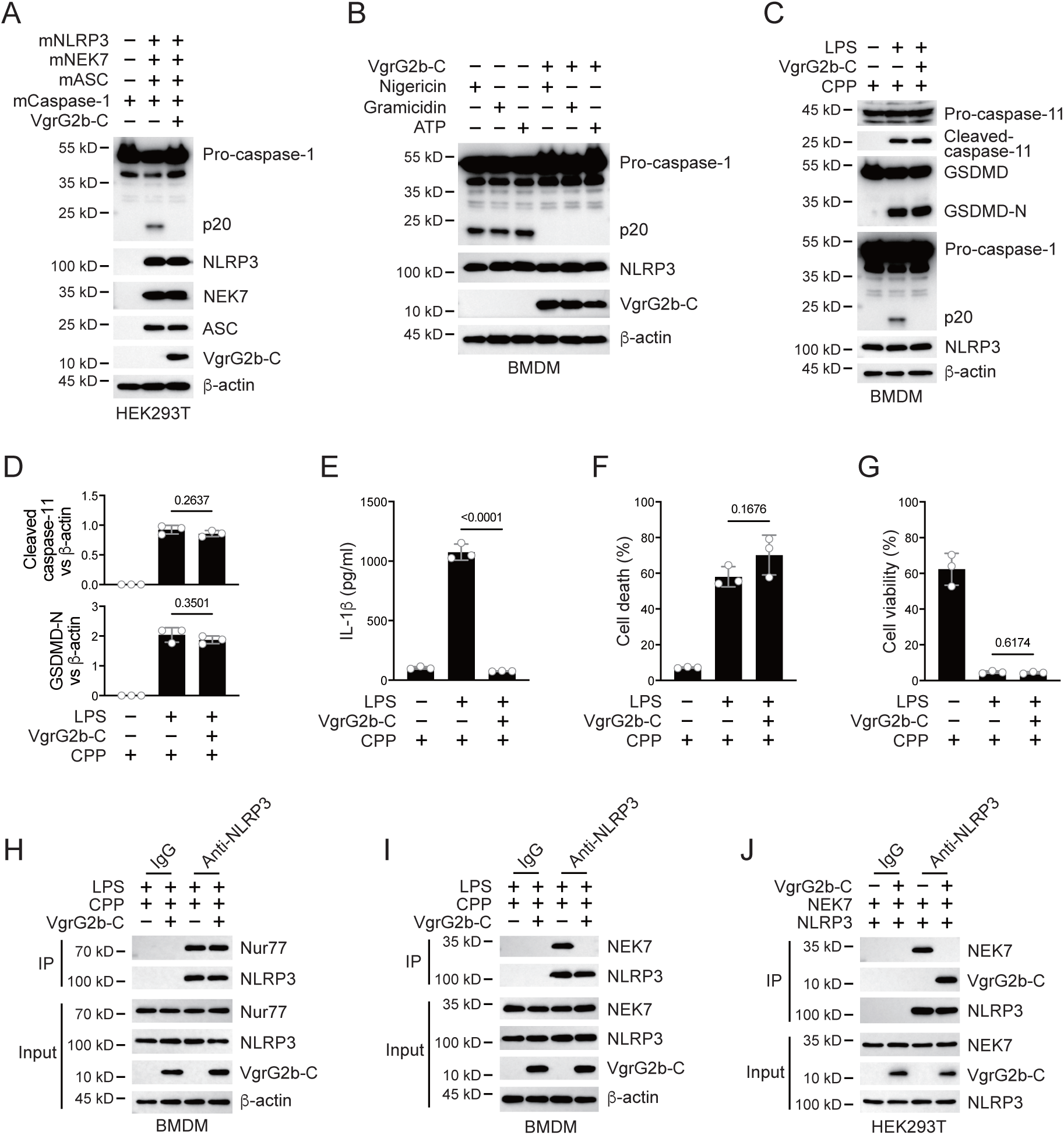
VgrG2b C-terminus suppresses the NLRP3 inflammasome. (A) Plasmids encoding VgrG2b C-terminus and the indicated murine proteins were co-transfected into HEK293T cells for 24 h, followed by immunoblotting with antibodies against the indicated proteins. (B) Wild-type BMDM cells were infected with lentiviruses encoding a control vector or VgrG2b C-terminus and primed with 1 μg/ml LPS for 3 h, followed by stimulation of 10 μM nigericin for 30 min, 0.5 μM gramicidin for 1 h and 2 mM ATP for 30 min. Cells were lysed and immunoblotted as indicated. (C–G) Wild-type BMDM cells were primed overnight with 1 μg/ml Pam3CSK4, followed by transfection of 2 μg/ml LPS using DOTAP with or without the transfection of VgrG2b C-terminus proteins using cell-penetrating peptides (CPP) for 2 h. Cells were then supplemented with fresh medium. Cells were lysed and immunoblotted 16 h post transfection (C). Band intensities of cleaved caspase-11 (top) and GSDMD (bottom) were quantified and compared to that of β-actin (D). Cell culture supernatants were collected for an ELISA assay to determine the secreted IL-1β protein levels 16 h post infection (E). Cytotoxicity was determined by LDH release assay in cell culture supernatants 16 h post infection (F). Cell viability was determined by an ATP quantification assay in cell pellets 16 h post infection (G). (H, I) Wild-type BMDM cells were primed overnight with 1 μg/ml Pam3CSK4, followed by transfection of 2 μg/ml LPS using DOTAP with or without the transfection of VgrG2b C-terminus proteins using cell-penetrating peptides (CPP) for 2 h. Cells were then supplemented with fresh medium. Cells were lysed and immunoprecipitated with a control IgG or antibody against NLRP3 16 h post transfection. Precipitates were immunoblotted with antibody against Nur77 (H) or NEK7 (I). (J) Plasmids encoding VgrG2b C-terminus, NEK7 and NLRP3 were co-transfected into HEK293T cells for 24 h, followed by immunoprecipitation with a control IgG or antibody against NLRP3. Precipitates were immunoblotted as indicated. Data were shown as means±SD. For 5E, 5F and 5G, data of three independent experiments were calculated. Experiments were repeated three times with similar results.

These data indicate that VgrG2b C-terminus exhibits a broad-spectrum inhibition of the NLRP3 inflammasome. Introducing VgrG2b C-terminus into BMDM cells through cell-penetrating peptides can significantly inhibit the activation of NLRP3 post intracellular LPS stimulation, without affecting the activation of caspase-11 and GSDMD (Fig. 5C, D). The secretion of IL-1β was significantly decreased (Fig. 5E). The cell death (Fig. 5F) and survival rates (Fig. 5G) were not affected. The activation of NLRP3 could also be inhibited by introducing VgrG2b C-terminus through cell-penetrating peptides when classical NLRP3 inflammasomes stimuli were present (Fig. S5C), accompanied by a reduction of IL-1β secretion (Fig. S5D), a decrease in cell death (Fig. S5E) and an increase in cell viability (Fig. S5F).

We next searched for the specific mechanism by which VgrG2b C-terminus inhibited the NLRP3 inflammasome. We first tested the ability of NLRP3 to catalyze ATP hydrolysis after the addition of VgrG2b C-terminus and found that the hydrolysis potential of NLRP3 was not affected (Fig. S5G). Meanwhile, VgrG2b C-terminus did not influence the binding of NLRP3 to mitochondrial DNA (Fig. S5H), as well as to Nur77 (Fig. 5H). The binding of NLRP3 to NEK7 was immensely inhibited by the C-terminus of VgrG2b (Fig. 5I). VgrG2b C-terminus did not bind to NEK7 (Fig. S5I). Domain mapping showed that the VgrG2b C-terminus bound to the same NLRP3 segment as NEK7 (Fig. 4G–I). We therefore detected the competitive binding between VgrG2b C-terminus and NEK7 to NLRP3. VgrG2b C-terminus competed against NEK7 (Fig. 5J), while NEK7 couldn’t compete with VgrG2b C-terminus (Fig. S5J), indicating that VgrG2b C-terminus has a stronger binding ability with NLRP3 compared to NEK7. Binding assays showed that VgrG2b-C had a tight binding ability for NLRP3 than NEK7 (326 nM VS 681 nM). Indeed, VgrG2b C-terminus did not inhibit the NLRC4 inflammasome (Fig. S5K).

### VgrG2b cleavage is essential for *P. aeruginosa* infection

Finally, we infected mice with a *P. aeruginosa* strain carrying a D883A mutation in the VgrG2b gene. The survival rate of mice was greatly improved when infected with the mutant strain (Fig. 6A). the level of IL-1β in the serum was also increased (Fig. 6B), while other inflammatory cytokines such as TNF-α and IL-6 showed little difference among wild-type and mutant bacteria (Fig. 6C, D). The bacterial load in the lungs of mice infected with the mutant strain was also decreased (Fig. 6E). We also detected the activation of inflammasomes in macrophages from lung lavage fluid and found that the activation of the NLRP3 inflammasome was activated in cells infected with the mutant strain (Fig. 6F), but the cell viability was not affected (Fig. 6G). We detected the protein levels of cytokines in the lavage fluid and found that the level of IL-1β was increased from mice infected with *P. aeruginosa* mutant strain (Fig. 6H). Other inflammatory cytokines such as TNF-α and IL-6 showed little difference (Fig. 6I, J).

**Figure 6.**
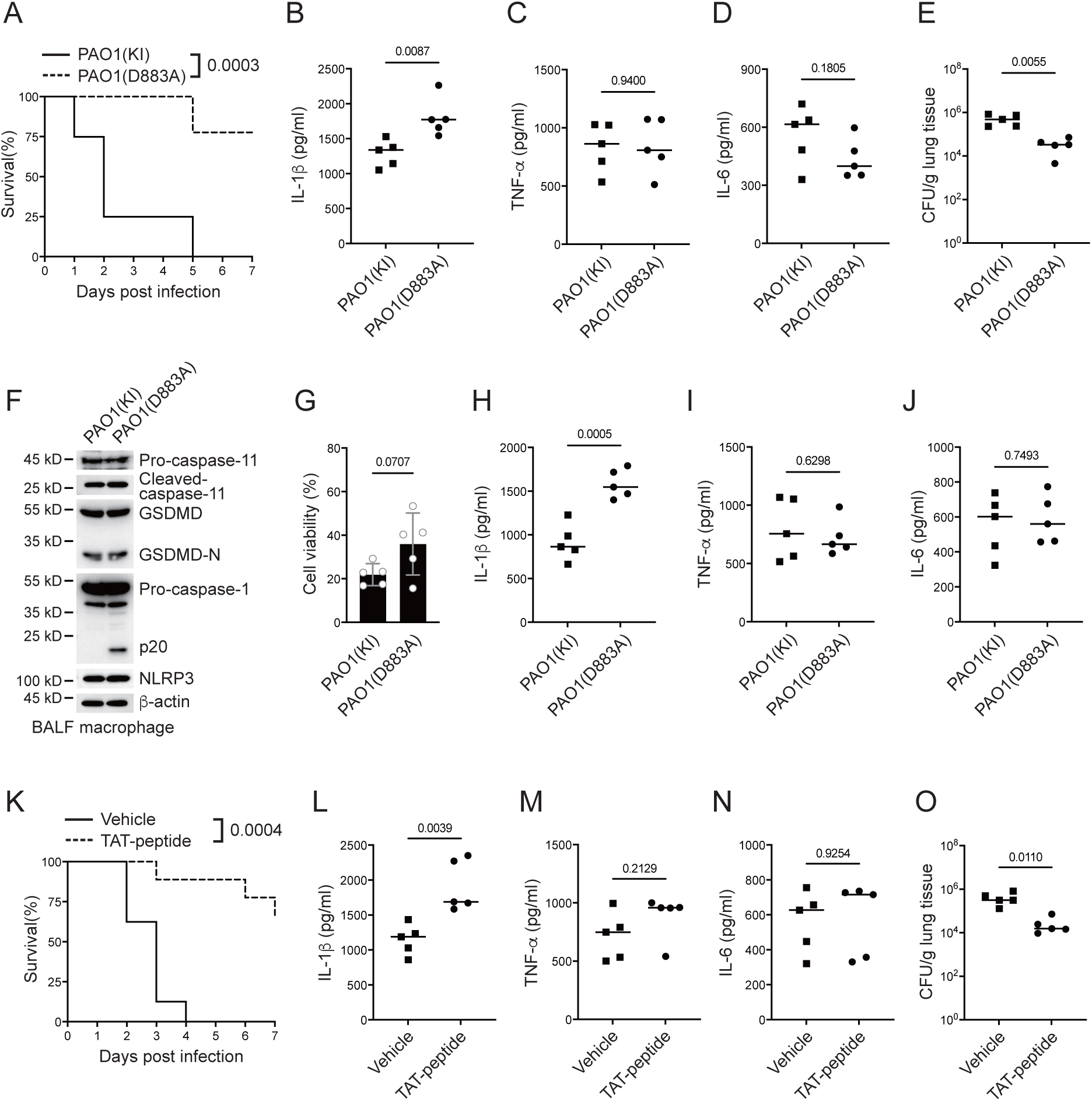
*P. aeruginosa* suppresses NLRP3 inflammasome *in vivo*. (A–E) Wild-type mice were intraperitoneally challenged with poly(I:C) at a dose of 2 mg/kg body weight for 6 h and then intranasally infected with 1x10^9^ cfu of VgrG2b (KI) or mutant (D883A) knockin ΔRetS PAO1, followed by survival calculation at the indicated days (A). Serum protein levels of IL-1β (B), TNF-α (C) and IL-6 (D) were examined through ELISA assays 24 h post infection. Bacterial load was examined in mouse lung tissues 4 days post infection (E). (F–J) Wild-type mice were intraperitoneally challenged with poly(I:C) at a dose of 2 mg/kg body weight for 6 h and then intranasally infected with 1x10^9^ cfu of VgrG2b (KI) or mutant (D883A) knockin ΔRetS PAO1. Bronchoalveolar lavage fluids (BALF) were collected 16 h post infection. Cells were immunoblotted as indicated (F). Cell viability was also determined by an ATP quantification assay in cell pellets (G). Protein levels of IL-1β (H), TNF-α (I) and IL-6 (J) in BALF supernatants were examined through ELISA assays. (K–O) Wild-type mice were intraperitoneally challenged with poly(I:C) at a dose of 2 mg/kg body weight for 6 h and then intranasally infected with 1x10^9^ cfu of ΔRetS PAO1 and administrated with 20 μg TAT sequence containing VgrG2b C-terminus peptide, followed by survival calculation at the indicated days (K). Serum protein levels of IL-1β (L), TNF-α (M) and IL-6 (N) were examined through ELISA assays 24 h post infection. Bacterial load was examined in mouse lung tissues 4 days post infection (O). Data were shown as means±SD. Experiments were repeated three times with similar results.

We designed a short peptide of VgrG2b C-terminus containing the VgrG2b linker region with a D883A substitution and a transmembrane TAT sequence. Peptides were then myristoylated (Zuber et al., 1989), aiming to target this peptide to inner membrane and suppress bacterial VgrG2b cleavage through competing with wild-type VgrG2b for caspase-11 (Fig. S6A). We first conducted tests at the cellular level and found that adding this peptide to BMDM cells after incubation with *P. aeruginosa* relieved the inhibition of the NLRP3 inflammasome (Fig. S6B) and the secretion of IL-1β was significantly increased (Fig. S6C). The cell death rate did not change significantly and the viability rate was not affected by this peptide (Fig. S6D, E). The same effect was observed in human macrophages, where this designed peptide could relieve the inhibition of *P. aeruginosa* on the NLRP3 inflammasome (Fig. S6F) and the secretion of cytokine IL-1β was also increased (Fig. S6G). Although it had no effect on cell death and viability (Fig. S6H, I), Finally, we delivered this peptide into the lungs of mice infected with *P. aeruginosa* through nebulization, and found that the survival rate of mice was increased (Fig. 6K). The level of IL-1β in the serum was increased (Fig. 6L), while other inflammatory factors such as TNF-α and IL-6 remained unchanged (Fig. 6M, N). The bacterial load was also decreased (Fig. 6O), indicating the potential medicinal value of this designed peptide in the future clinical treatment of *P. aeruginosa* infections.

## Discussion

In this study, we found that the structural protein VgrG2b from *P. aeruginosa* can be cleaved by caspase-11 in host cells. The C-terminal fragment of VgrG2b bound to NLRP3, inhibiting the activation of the NLRP3 inflammatory and the secretion of IL-1β. The delivery of VgrC2b C-terminus mimics significantly improved the infection symptoms caused by *P. aeruginosa*, providing an optional treatment approach for chronic infections of *P. aeruginosa*.

The infection of *P. aeruginosa*, especially those in chronic infections, is an important factor causing pulmonary fibrosis in clinicals (Qin *et al*., 2022; Rossi *et al*., 2021). LPS from Gram-negative bacterium can be transported to host cells and activate non-canonical inflammasomes within them (Kayagaki *et al*., 2013; Shi *et al*., 2014). There are many ways for LPS from bacteria to be transported into host cells, including binding with HMGB1 originated from the liver and entering the interior of the cells through the surface of macrophages, as well as enveloping and delivering LPS into the host cells through extracellular vesicles (Deng et al., 2018; Vanaja et al., 2016). We observed the activation of non-canonical inflammasomes in macrophages after infection with *P. aeruginosa*, indicating that LPS was also transported into the cells. *P. aeruginosa* is not a typical intracellular bacterium, and its LPS entry into target cells is likely through the out membrane vesicles *in vivo*.

*P. aeruginosa* mainly utilizes the T6SS secretion system to promote the establishment of chronic infections (Qin *et al*., 2022). During this process, H2-T6SS plays many important roles, responsible for delivering known and yet to be explored virulence factors into host cells (Jiang *et al*., 2014; Sana *et al*., 2015). Here we have found that VgrG2b is not only responsible for the delivery of bacterial virulence proteins, but it can also serve as a virulence factor on its own. Its uniqueness lies in the fact that it requires cutting to be effective. Whether there are other proteins secreted through the H2-T6SS in *P. aeruginosa* that need to be cleaved by host proteases before they exert their function is a very worthwhile research topic. Of course, this processing from the host is not limited to cleavage, but may also be post-translational modifications of proteins such as phosphorylation or ubiquitination (Ashida et al., 2014; Li et al., 2021). Systematically identifying these possible modifications and their substrates will be of great significance for understanding the pathogenic mechanism of *P. aeruginosa*.

The non-canonical inflammasome receptor caspase-11 has a very narrow substrate spectrum (Shi *et al*., 2023). The process of substrate cleavage by inflammatory caspases can be divided into two parts. Firstly, inflammatory caspases bind to a part of the substrate, which is often not their cleavage site, and the other is the recognition of the true site of their cleavage. This mechanism ensures that limited proteins are cleaved by host cells when inflammasomes are activated, thereby confining the severity of the inflammatory response. Here, we found that caspase-11/4 specifically cleaves the structural proteins of the secretion system of *P. aeruginosa*, and the generated new fragments can play an inhibitory role. This is a new way for pathogenic microorganisms to hijack the host’s immune system. Because traditional hijacking usually directly affects the host immune system without cutting. It is very effective for *P. aeruginosa* using VgrG2b fragment to inhibit NLRP3. On the one hand, VgrG2b transports virulence factors into the host cell, and on the other hand, the cleaved substrate plays a role in inhibiting the NLRP3 inflammasome.

In summary, we have discovered a new pattern of recognition between bacterial proteins and the host immune system during *P. aeruginosa* infection, which involves hijacking the host’s protease to produce a new virulence factor that can inhibit the host’s immune response.

### Experimental procedures

#### Antibodies and reagents

Antibodies against NLRP3 (15101), IL-1β (12703, 12426), cleaved IL-1β (83186, 63124), GSDMD (93709), caspase-1 (3866, 24232, 2225) and ASC (13833, 67824) were purchased from Cell Signaling Technology. Antibodies against caspase-1 (ab207802), NEK7 (ab133514), Nur77 (ab1533914) and GSDMD (ab215203) were purchased from Abcam. Antibodies against caspase-11 were: ab180673 from Abcam, 14340 from Cell Signaling Technology, 14-9935-82 from Invitrogen, M029-3 from MBL Beijing Biotech. Anti-c-Myc magnetic beads (88843) were purchased from Pierce. Antibody against 6xHis tag (MA121315) was purchased from Invitrogen. Antibodies against FLAG tag (F3165), β-actin (A1978) and anti-FLAG M2 magnetic beads (M8823) were purchased from Sigma-Aldrich. Antibodies against c-Myc (sc-40), Tom20 (sc-11415), GAPDH (sc-32233) and GST (sc-138) were purchased from Santa Cruz Biotechnology. Anti-HA tag antibody (HX1820) was purchased from Huaxingbio (Beijing). HRP-conjugated goat anti-mouse IgG (SA00001-1) and HRP-conjugated goat anti-rabbit IgG (SA00001-2) were purchased from Proteintech Group. Alexa Fluor Plus 488-conjugated goat anti-mouse IgG (A32723) and Alexa Fluor 594-conjugated goat anti-rabbit IgG (A32740) were purchased from Invitrogen. Alexa Fluor Plus 488-conjugated donkey anti-goat IgG (bs-0294D-AF488) and Alexa Fluor 594-conjugated donkey anti-rabbit IgG (bs-0295D-AF594) were purchased from Bioss (Beijing).

Glutathione sepharose 4B resin (17075601) was purchased from Cytiva. Protein A/G PLUS-Agarose beads (sc-2003) were purchased from Santa Cruz Biotechnology. DAPI (2879038) was purchased from PeproTech (BioGems). SMART MMLV reverse transcriptase (639524) was purchased from Clontech, Takara Bio. Random primer (Hexadeoxyribonucleotide mixture, pd(N)6) (3801), X-α-Gal (630462) and carrier DNA (630440) were purchased from Takara Bio. Protease inhibitor cocktail (11697498001) was purchased from Roche. Total RNA extraction kit (8034111) was purchased from Dakewe (Beijing). Ni-NTA agarose beads (R90115) and AminoLink coupling resin (20381) were purchased from Invitrogen. LPS (B46894) was purchased from Innochem (Beijing). DOTAP was: D10530 from Psaitong (Beijing) and 11202375001 from Roche. Nigericin (481990), Gramicidin (368020), ATP (A6559) were purchased from Merck Millipore. Poly(I:C) (tlrl-picw) and Pam3CSK4 (tlrl-pms) were purchased from InvivoGen. Cell-penetrating peptides: ab142343 from Abcam and sc-396807 from Santa Cruz. Human M-CSF (300-25) and murine M-CSF (315-02) were purchased from PeproTech. CellTiter-Glo luminescent cell viability assay kit (G7570) and CytoTox 96 non-radioactive cytotoxicity assay kit (G1780) were purchased from Promega.

### Bacteria and CFU determination

*E. coli* (CGMCC1.2389, ATCC11775) was from China General Microbiological Culture Collection Center. *Pseudomonas aeruginosa* (PAO1) was a gift from Dr. Lvyan Ma (Institute of Microbiology, Chinese Academy of Sciences). *E. coli* and *P. aeruginosa* were preserved in LB plates and inoculated to liquid LB medium, followed by shaking at 37 °C until they grew to the exponential phase (optical density=0.6–0.8). Bacteria were diluted and spread into LB plates to determine the CFU numbers. When the bacteria were cultured to the mid-exponential stage, they were washed three times with PBS and resuspended in PBS or DMEM.

### *P. aeruginosa* OMV purification

When *P. aeruginosa* was cultured to the mid exponential stage, cultures were centrifugated to obtain the supernatant. After filtering through a 0.22 μm filter, the supernatant was placed in a discontinuous sucrose gradient tube and centrifuged at a speed of 100, 000 g for 16 h. The precipitate was resuspended by PBS and centrifuged again to remove residual culture medium components. The precipitate was resuspended with PBS again, resulting in the OMV sample. The sample was immediately used or frozen and stored in a –80 °C freezer. 20 μg/ml OMV was used in each *in vitro* experiment.

### *P. aeruginosa* knockout and knockin strategy

The knockout plasmid pK18mobsacBGm was digested with BamHI and HindIII, and then recovered by agarose gel electrophoresis. Primers were designed to clone homologous arms of approximately 900 bp upstream and downstream of the target gene. This knockout strategy preserved the starting 99bp and ending 99bp of the target gene. The upstream and downstream homologous arms were then connected with the linearized pk18mobsacBGm vector using homologous recombination, and the knockout vector pk18mobsacBGm-target were obtained through sequencing. Primers used were: RetS-up-F: 5’-ctcggtacccggggatccccgccgtgcgcgacatgctcgccggcaa-3’, RetS-up-R: 5’-agcacgtcgctgc ccggcgaagtcccttcg-3’, RetS-down-F: 5’-ccttcgaagggacttcgccgggcagcgacgtgctccgg-3’, RetS-down-R: 5’-cgttgtaaaacgacggccagtgccaagcttcgagggtcaggcaggcgag-3’ for RetS knockout; KO-VgrG2b-up-F: 5’-attcgagctcggtacccggggatcccttgatggaaaagagtttcaagacc-3’, KO-VgrG2b-up-R: 5’-tctcgaggaaataatctcgaacgataggctcgcagagcgcttcttccagt-3’, KO-VgrG2b-down-F: 5’-actggaagaagcgctctgcgagcctatcgttcgagattatttcctcga-3’, KO-VgrG2b-down-R: 5’-taaaacgacggccagtgccaagcttgacgacgtcggggttctctgcctt-3’ for VgrG2b knockout; KI-Myc-VgrG2b-up-F: 5’-tcgagctcggtacccggggatccgagcacatcaccctgatgtgcggcggcgcct-3’, KI-Myc-VgrG2b-up-R1: 5’-ttcgaaccgcgggccctctagactcgagcggtatcccgttgggaagtttttcagt-3’, KI-Myc-VgrG2b-up-R2: 5’-atcctcttctgagatgagtttttgttcgaaccgcgggccctctagactc-3’, KI-Myc-VgrG2b-down-F: 5’-aaaaactcatctcagaagaggatctgtgaccaatgaaatgcaagaccttgctca-3’, KI-Myc-VgrG2b-down-R: 5’-taaaacgacggccagtgccaagcttcggccccgccggcgccactggcgaa-3’ for VgrG2b-Myc knockin; VgrG2b-D883A-up-F: 5’-aattcgagctcggtacccggggatccgagcaccagg gcgtggggcacgacg-3’, VgrG2b-D883A-up-R: 5’-aggtcttgcatttcattggtcacagatcctcttctgagat gag-3’, VgrG2b-D883A-down-F: 5’-actcatctcagaagaggatctgtgaccaatgaaatgcaagaccttgctc a-3’, VgrG2b-D883A-down-R: 5’-taaaacgacggccagtgccaagcttttcgccggccaggcagaattcgacg-3’ for VgrG2b D883A knockin. The recipient bacterium *P. aeruginosa*, co-plasmid pRK2013, and donor bacterium (pK18mobsacBGm-target) were inoculated in LB liquid medium (with corresponding antibiotics added), and incubated overnight in a shaking table at 37 °C and 200 rpm. 1 ml of co-plasmid and donor bacterial culture medium were collected separately and centrifuged at 5, 000 rpm for 2 m. Pellets were collected and resuspended in 200 μl LB medium. Co-plasmids and donor bacteria were mixed in a 1:1 ratio, and inoculated on LB solid culture medium at 37 °C for 2 hours. Simultaneously, the recipient strain *P. aeruginosa* was placed in 42 °C for 2 hours of cultivation. 1mL of receptor bacteria was collected and centrifuged at 5, 000 rpm for 2 min and resuspended in 200 μl LB liquid medium. The bacteria were spotted onto a mixed plaque of co-plasmids and donor bacteria and cultured at 37 °C for 4 hours. The plaque was picked up and suspended in the 500–800 μl LB liquid medium and plated onto LB plates with Irg and Gm resistance, followed by culture overnight at 37 °C. A single colony was selected and transferred again to LB plates with Irg and Gm resistance. The next day, bacteria were drawn a line on a 20–22% sucrose agar plate and cultured at 37 °C for 24–48 hours. Single colonies were selected again according to the above method. Colonies that could grow on LB plates but not on Gm resistant plates were chosen and subjected to PCR validation.

### Cell component fractionation and mitochondrial DNA quantification

Cells were fractionated according to the guidelines of a cell fractionation kit from Abcam (ab109719). Mitochondria were isolated from macrophages using a mitochondrial isolation kit from Abcam (ab110170) or from Invitrogen (89874), followed by mitochondrial DNA purification through a mitochondrial DNA isolation kit from Abcam (ab65321). Total cytosolic DNA was extracted using a DNeasy Blood & Tissue Kit (QIAGEN) following manufacturer’s instructions. For mitochondrial DNA quantification, samples containing mitochondrial DNA were subjected to PCR analyses using the following primers: 5’-GAGATGACCAAATTTACAATG-3’ (forward) and 5’-TCCTGTTCCTGCTCCTGCTTC-3’ (reverse) for mouse mitochondrial gene COX1.

### Mice and infection

*Casp4*^−/−^ (S-KO-01332), *Gsdmd*^−/−^ (S-KO-12963) and *Aim2*^−/−^ (S-KO-09889) mice were purchased from Cyagen Biosciences (Jiangsu, China). Mice were intraperitoneally challenged with poly(I:C) at a dose of 2 mg/kg body weight for 6 h and then intranasally infected with 1x10^9^ cfu of *P. aeruginosa* strain PAO1, followed by survival calculation at the indicated days and bacterial load determination 4 days later. Sera were collected and subjected to ELISA assays. For bronchoalveolar lavage fluid (BALF) collection, mice were anaesthetized, followed by thoracic cavity opened and a flush of circulating blood cells by injecting about 10 ml ice-cold PBS into the right ventricle. Mouse neck was then dissected to expose trachea. A puncture needle was inserted into the upper end of the trachea, followed by repeated rinsing with 0.8 ml PBS for five times. BALF collections were centrifuged at 300 g at 4 °C for 5 min, followed by ELISA assays of the supernatants or macrophage isolation in the cell pellets.

### Primary cell separation and culture

To generate BMDM cells, bone marrow cells were flushed from mouse femurs and tibias, followed by red blood cell removal using red blood cell lysis buffer. Cells were resuspended in alpha-MEM containing 10% (v/v) fetal bovine serum and 10 ng/ml M-CSF and grown at 37 °C with a 5% CO_2_ humidified atmosphere for 7 days. To produce human macrophages, human bloods collected from healthy donors were mixed with an equal volume of saline and then added to lymphocyte separation medium, followed by centrifugation and PBMC collection. Human PBMCs were incubated with APC-conjugated anti-CD14 antibody on ice for 30 min, followed by incubation with anti-APC-conjugated magnetic beads. Monocytes were enriched on a magnetic separator. To test the purity of sorted monocytes, cells were stained with APC conjugated anti-CD14 antibody, followed by FACS examination. Cells with purity above 95% were used for subsequent experiments. Cells were resuspended in RPMI-1640 medium containing 10% (v/v) fetal bovine serum and 50 ng/ml human M-CSF, followed by growing at 37 °C with a 5% CO_2_ humidified atmosphere for 7 days. Healthy blood samples were provided by the researchers volunteering. Informed consents were obtained from all subjects and experiments conformed to related principles. Study was licensed by the Ethics Committee of Institute of Microbiology, Chinese Academy of Sciences.

### Inflammasome induction

For canonical NLRP3 inflammasome activation, cells were incubated with 1 μg/ml LPS for 3 h, followed by stimulation with 10 μM nigericin for 30 min, 0.5 μM gramicidin for 1 h and 2 mM ATP for 30 min. Cells were grown at 37 °C with a 5% CO_2_ humidified atmosphere. For non-canonical inflammasome stimulation, macrophages were primed overnight with 1 μg/ml Pam3CSK4, followed by transfection of 2 μg/ml LPS using DOTAP for 16 h. Otherwise, activation induced by *P. aeruginosa* infection, bacteria were cultured in LB at 37 °C until OD_600_ reached 0.8, then washed by PBS and diluted in serum-free DMEM medium to infect cells. Macrophages were primed overnight with 1 μg/ml Pam3CSK4 and incubated with *P. aeruginosa* strain PAO1 at an MOI of 30 and OMVs for 2 h. Cells were then supplemented with fresh medium containing 100 μg/ml Gentamycin and cultured for 16 h.

### Plasmid transfection and lentiviral production

Transfection reagent JetPRIME (Polyplus Transfection) was used to conduct transfection in HEK293T cells. HEK293T cells were plated 14 h earlier to achieve a density of about 70% when transfection was performed. Lentiviral plasmids containing exogenous genes of interest and packaging plasmids pSPAX2 and pMD2.G were transfected into HEK293T cells at a ratio of 4:3:1. 48 h later, supernatants were collected, followed by centrifugation at 1,000 rpm for 5 min and filtration using 0.45 μm sterile syringe filters (Merck Millipore). The filtrate was then concentrated with a 100 kD Amicon ultra centrifugal filter unit (Merck Millipore). Macrophage cells were incubated with lentiviruses for 6 h, followed by culture medium replacement with fresh medium. 2 μg/ml puromycin was added to the medium 48 h post infection for 3 days.

### Protein transfection using cell-penetrating peptides (CPP)

For a single well of 6-well plate macrophages, 10 μg of wild-type or mutant VgrG2b-C (884-1019) was incubated with 10 μg cell-penetrating peptides in 100 μl of PBS at room temperature for 30 min, followed by addition of the peptide-CPP mixture to the cell medium. The culture medium was replaced with fresh medium later.

### Cell viability and cytotoxicity assay

Cells following inflammasome activation were centrifuged at 187 g for 5 min, and supernatants and cell pellets were collected, respectively. Cell pellets from cells stimulated with inflammasome activators were used to measure cell viability through checking cellular ATP levels with a CellTiter-Glo luminescent cell viability assay kit (Promega). The supernatants from cells stimulated with inflammasome activators were collected and subjected to lactate dehydrogenase (LDH) assay to assess cell death using a CytoTox 96 non-radioactive cytotoxicity assay kit (Promega) and cellular cytotoxicity was displayed as a percentage of total cellular LDH (100% cell lysis control). Both the supernatants and cell pellets were collected for immunoblotting analyses.

### ELISA

ELISA kits detecting human IL-1β (D711068), mouse IL-1β (D721017), mouse IL-6 (D721022) and mouse TNF-α (D721217) were purchased from BBI life sciences (Shanghai). Cell culture supernatants were collected and subjected to ELISA assays following manufacturer’s instructions.

### RNA extraction and RT-PCR

A total RNA extraction kit (Dakewe, Beijing) was used to extract RNA from cells according to the manufacturer’s instructions. For reverse transcription, 1 μg RNA was mixed with 2 μl of N6 primers and 7 μl of DEPC water, followed by incubation at 72 °C for 2 min. The mixture was then chilled on ice for 2 min and mixed with 4 μl of 5x first-strand buffer (TaKaRa Bio), 2 μl of 100 mM DTT, 2 μl of 10 mM dNTP mix (Solarbio, Beijing) and 2 μl of SMART MMLV reverse transcriptase (Clontech, TaKaRa Bio). The mixture was incubated under the following conditions: at 25 °C for 10 min, at 42 °C for 1 h and at 75 °C for 10 min. cDNA samples were diluted for further PCR analysis.

### Recombinant protein expression and purification

For recombinant expression of PAO1 VgrG2b (full-length and C-terminus), cDNA of full-length PAO1 VgrG2b, amino acids (aa) 884-1019, mutant (H935A;H936A;H939A;E983A) aa 884-1019 was cloned into pGEX-6P-1 vector with an N terminal GST tag and a C terminal Myc tag. Plasmids were transformed into *E. coli* strain BL21 and the bacteria grew in LB medium supplemented with 100 μg/ml ampicillin at 37 °C for 8 h, followed by addition of IPTG to a final concentration of 0.5 mM after OD_600_ reached 0.8 and shaking at 150 rpm at 37 °C for 3 h. Cells were harvested and resuspended with PBS containing 1% Triton-X-100 and lysed by sonication. Cell lysates were centrifuged at 13,000 rpm at 4 °C for 30 min and filtered through sterile syringe filters. For crude purification of recombinant proteins, the filtered lysate was loaded on column containing glutathione sepharose 4B resin, followed by washing with PBS containing 1% Triton-X-100. The protein was eventually eluted by 1 ml 10 mM L-Glutathione solution. For removal of LPS contaminations, crudely purified proteins were further eluted through Pierce high-capacity endotoxin removal spin columns and heparin columns to remove LPS and bacterial genomic DNA, respectively. GST tags were removed by a PreScission protease.

For recombinant expression of caspase-11 p10 and p22, cDNAs encoding proteins aforementioned were cloned into pET-15b with a C-terminal 6xHis tag. Plasmids were transformed into *E. coli* strain BL21, and the bacteria grew in LB medium supplemented with 100 μg/ml ampicillin for 8 h at 37 °C. Protein expression was induced by 0.1 mM isopropyl β-D-1-thiogalactopyranoside (IPTG) through shaking at 120 rpm overnight at 16 °C. The whole purification procedure was performed at 4 °C. Cells were collected and resuspended in a binding buffer containing 0.5 M NaCl, 20 mM Tris, 5 mM imidazole, and 1% Triton X-100, pH 7.5, and lysed by sonication. The lysate was centrifuged at 16,200 g for 30 min. Cleared lysate was filtered through sterile syringe filters before passing through a column loaded with Ni-NTA agarose beads. The column was washed with a wash buffer containing 0.5 M NaCl, 20 mM Tris, and 60 mM imidazole, pH 7.5, and eluted by an elution buffer containing 1 M NaCl, 40 mM Tris, and 2 M imidazole, pH 7.5.

### *In vitro* cleavage assay

*In vitro* caspase cleavage was performed as previously described (Wang K, Cell, 2020). Hydrolysis of VgrG2b substrates by recombinant caspases were performed in a buffer containing 50 mM HEPES (pH 7.5), 150 mM NaCl, 3 mM EDTA and 0.005% (v/v) CHAPS, and 10 mM DTT. 1 μg recombinant VgrG2b protein with or without inhibitor were incubated with 0.5 mM caspase-11 p22/p10 complex in a 40 μl reaction buffer at 37 °C for 4 h. The mixture was added with SDS loading buffer and boiled for 10 min, followed by standard immunoblotting analysis.

### Immunoprecipitation

HEK293T cells were seeded on 6-well culture plates at a confluency of 70% and were transfected with plasmids 16 h later. 24 h post transfection, cells were washed twice with PBS and lysed in a pre-chilled buffer containing 150 mM NaCl, 50 mM Tris (pH 7.5), 1 mM EDTA, 0.5% digitonin, 1% protease inhibitor cocktail and 10% glycerol on ice for 30 min. For immunoprecipitation using primary cells, cells post stimulation or treatments were lysed in a pre-chilled PBS buffer containing 0.5% digitonin, 1% protease inhibitor cocktail on ice for 30 min. Cell lysates were centrifuged at 13,000 rpm at 4 °C for 10 min. Primary antibodies and their isotype control IgG were immobilized to AminoLink Coupling Resin (ThermoFisher Scientific) following the manufacturer’s instructions. Supernatants were incubated with immobilized antibodies at 4 °C for 2 h. Resins were washed three times with PBS containing 0.5% digitonin, followed by immunoprecipitant detachment in 0.1 M Glycine-HCl, pH 2.7. Supernatants were neutralized by adding 1/10 the volume of 2 M Tris-HCl, pH 8.0, followed by immunoblotting with the indicated antibodies.

### Yeast two-hybrid screening

Yeast strain AH109 was inoculated into 5 ml YPDA medium and shook at 250 rpm for 16 h at 30 °C. The overnight culture was transferred to a flask containing 50 ml YPDA and incubated for 3 h at 30 °C. Yeast cells were collected by centrifugation at 3,000 g for 10 min and resuspended in a 1X TE/LiAc buffer (pH7.5) containing 0.1 M lithium acetate, 0.1 M Tris–HCl, and 10 mM EDTA. A mixture of plasmid DNA and carrier DNA was added to the yeast cell suspension and vortexed to mix well. A PEG/LiAc buffer containing 40% polyethylene glycol 3350, 0.1 M lithium acetate, 0.1 M Tris–HCl, and 10 mM EDTA was then added to the suspension and vortexed at a high speed. The suspension was shaken at 200 rpm for 30 min at 30 °C, followed by addition of DMSO and heat shock for 15 min in a 42 °C water bath. Cells were chilled in an ice bath for 2 min and resuspended to plate on appropriate SD agar plates. A mouse bone marrow cDNA library was used for Y2H screening with BD-tagged VgrG2b-C as bait. BD-tagged VgrG2b-C truncation and AD-tagged NLRP3, NLRP6, NLRC4 were co-transfected into yeast strain AH109, and double-positive clones were grown on plates containing X-α-gal. The intensity of blue color was monitored on a scanner.

### Immunofluorescence

Cells adhered to covers pre-coated with poly-L-lysine were stimulated with inflammasome activators, followed by fixation with 4% paraformaldehyde for 10 min at room temperature. Cells were then washed twice with PBS and permeabilized with PBS containing 0.1% digitonin for 10 min, followed by washing with PBS for twice and blockage in 10% normal goat serum at 37 °C for 30 min. Cells were incubated with primary antibody for 2 h, followed by washing with PBS for three times and further incubation with fluorescence-conjugated secondary antibody for 1 h. Nuclei were stained with DAPI. Cells were visualized through an UltraVIEW VoX imaging system (PerkinElmer).

### Statistical analysis

No statistical methods were used to predetermine sample size. Experiments were independently repeated at least three times to achieve statistical significance. No randomization or blinding procedures was used in this study. No samples were excluded from the analysis. Data were shown as means ± SD of three technical replicates. Data with normal distribution determined by Shapiro–Wilk normality test were statistically analyzed by two tailed Student’s t tests if not specified. The Gehan-Breslow-Wilcoxon test was used for the analysis of survival data. Data were analyzed by GraphPad Prism 9.0. *P*-values ≤0.05 were termed as significant; NS means non-significant where *P*>0.05.

## Supporting information

Support Figure 1

Support Figure 2

Support Figure 3

Support Figure 4

Support Figure 5

Support Figure 6

## Acknowledgements

We thank Yingyu Chen (Peking University) for technical help. This work was supported by the National Natural Science Foundation of China (92369104, 82271790, 92169113), Beijing Natural Science Foundation (JQ23028, 7212067), the National Key R&D Program of China (2021YFA1300202, 2022YFC2302900), Strategic Priority Research Programs of the Chinese Academy of Sciences (XDB29020000), Key Research Program of Frontier Sciences of Chinese Academy of Sciences (ZDBS-LY-SM025), CAS Project for Young Scientists in Basic Research (YSBR-010), Fok Ying Tung Education Foundation to P.X., Youth Innovation Promotion Association of CAS to S.W.

## Author contributions

Y.Q. and Q.L. performed experiments and analyzed data; X.C., C.W., C.K., M.L., C.R., D.J. and S.W. performed experiments; P.X. initiated the study, designed and performed experiments, analyzed data, and wrote the paper.

