## Supplementary Figure Legends for "A VgrG2b fragment cleaved by caspase-11/4 promotes *Pseudomonas aeruginosa* infection through suppressing the NLRP3 inflammasome"

**Supplementary Figure 2. Caspase-11 cleaves limited VgrG family members.** (A, B) Plasmids encoding VgrG1a and mouse caspase-11 p22/p10 (A) or human caspase-4 p22/p10 (B) were co-transfected into HEK293T cells for 24 h, followed by immunoblotting with antibodies against the indicated proteins. (C–L) Plasmids encoding the indicated VgrG family members and mouse caspase-11 p22/p10 or human caspase-4 p22/p10 were co-transfected into HEK293T cells for 24 h, followed by immunoblotting with antibodies against the indicated proteins. (M) Wild-type BMDM cells were primed overnight with 1 μg/ml Pam3CSK4, followed by incubation of VgrG2b-Myc knockin (KI) or knockout (KO) ΔRetS PAO1 at an MOI of 30 and 20 μg/ml OMVs for 2 h. Cells were then supplemented with fresh medium containing 100 μg/ml Gentamycin. Cells were lysed and immunoblotted 16 h post infection. Band intensities of cleaved caspase-11 (top) and GSDMD (bottom) were quantified and compared to that of β-actin. Data were shown as means±SD. Experiments were repeated three times with similar results.

**Supplementary Figure 3. Caspase-11 cleaves VgrG2b during *P. aeruginosa* infection.** (A) Plasmids encoding VgrG2b and mutant caspase-4 p22/p10 were co-transfected into HEK293T cells for 24 h, followed by immunoprecipitation with a control IgG or antibody against Myc. Precipitates were immunoblotted as indicated. (B) Scheme for VgrG2b chimeras. (C) Plasmids encoding VgrG2b-N-GFP chimera and caspase-11 p22/p10 were co-transfected into HEK293T cells for 24 h, followed by immunoblotting with antibodies against the indicated proteins. (D) Plasmids encoding VgrG2b-C and mutant caspase-4 p22/p10 were co-transfected into HEK293T cells for 24 h, followed by immunoprecipitation with a control IgG or antibody against Myc. Precipitates were immunoblotted as indicated. (E) Recombinant VgrG2b was incubated with caspase-11 p22/p10 subunits with or without the presence of 10 μM Ac-FLTD-CMK, followed by immunoblotting with antibodies against the indicated proteins. (F) BMDM cells were primed overnight with 1 μg/ml Pam3CSK4, followed by transfection of 2 μg/ml LPS using DOTAP with or without the incubation of VgrG2b-Myc knockin (KI) ΔRetS PAO1 at an MOI of 30 for 2 h. Cells were then supplemented with fresh medium containing 100 μg/ml Gentamycin and 10 μM Ac-FLTD-CMK or 10 μM z-DEVD-FMK. Cells were lysed and immunoblotted 16 h post infection. (G–K) Human macrophages were primed overnight with 1 μg/ml Pam3CSK4, followed by transfection of 2 μg/ml LPS using DOTAP with or without the incubation of VgrG2b (KI) or mutant (D883A) knockin ΔRetS PAO1 at an MOI of 30 for 2 h. Cells were then supplemented with fresh medium containing 100 μg/ml Gentamycin. Cells were lysed and immunoblotted as indicated 16 h post infection (G). Band intensities of cleaved caspase-4 (top) and GSDMD (bottom) were quantified and compared to that of β-actin (H). Cell culture supernatants were collected for an ELISA assay to determine the secreted IL-1β protein levels 16 h post infection (I). Cytotoxicity was determined by LDH release assay in cell culture supernatants 16 h post infection (J). Cell viability was determined by an ATP quantification assay in cell pellets 16 h post infection (K). Data were shown as means±SD. For S3I, S3J and S3K, data of three independent experiments were calculated. Experiments were repeated three times with similar results.

**Supplementary Figure 4. Cleaved VgrG2b fragment binds to NLRP3.** (A) Plasmids encoding VgrG2b N-terminus and NLRP3 were co-transfected into HEK293T cells for 24 h, followed by immunoprecipitation with a control IgG or antibody against Myc. Precipitates were immunoblotted as indicated. (B) VgrG2b C-terminus proteins were introduced into human macrophage cells with the help of cell-penetrating peptides (CPP) for 6 h. Cells were lysed and immunoprecipitated with a control IgG or antibody against Myc. Precipitates were immunoblotted as indicated. (C) Human macrophage cells were primed overnight with 1 μg/ml Pam3CSK4, followed by transfection of 2 μg/ml LPS using DOTAP with the incubation of VgrG2b knockin (KI) ΔRetS PAO1 at an MOI of 30 for 2 h. Cells were then supplemented with fresh medium containing 100 μg/ml Gentamycin. Cells were lysed and immunoprecipitated with a control IgG or antibody against NLRP3 16 h post infection. Precipitates were immunoblotted as indicated. (D–G) Plasmids encoding VgrG2b C-terminus and GSDMD (D), ASC (E), caspase-1 (F) and caspase-1 p20/p10(G) were co-transfected into HEK293T cells for 24 h, followed by immunoprecipitation with a control IgG or antibody against Myc. Precipitates were immunoblotted as indicated. (H) *Gsdmd*^+/+^ and *Gsdmd*^–/–^ BMDM cells were primed overnight with 1 μg/ml Pam3CSK4, followed by transfection of 2 μg/ml LPS using DOTAP with the incubation of VgrG2b-Myc knockin (KI) ΔRetS PAO1 at an MOI of 30 for 2 h. Cells were then supplemented with fresh medium containing 100 μg/ml Gentamycin. Cells were lysed and immunoprecipitated with a control IgG or antibody against NLRP3 16 h post infection. (I) Plasmids encoding wild-type VgrG2b, mutant VgrG2b (H935A;H936A;H939A;E983A) and caspase-11 p22/p10 were co-transfected into HEK293T cells for 24 h, followed by immunoblotting with antibodies against the indicated proteins. (J) Plasmids encoding wild-type VgrG2b C-terminus, mutant VgrG2b C-terminus (H935A;H936A;H939A;E983A) and NLRP3 were co-transfected into HEK293T cells for 24 h, followed by immunoprecipitation with a control IgG or antibody against Myc. Precipitates were immunoblotted as indicated. (K) BMDM cells were primed overnight with 1 μg/ml Pam3CSK4. Recombinant wild-type VgrG2b C-terminus and mutant VgrG2b C-terminus (H935A;H936A;H939A;E983A) were introduced into BMDM cells with the help of cell-penetrating peptides (CPP) for 6 h, followed by transfection of 2 μg/ml LPS using DOTAP. Cells were immunoblotted with antibodies against the indicated proteins. Precipitates were immunoblotted as indicated. Data were shown as means±SD. Experiments were repeated three times with similar results.

**Supplementary Figure 5. VgrG2b C-terminus inhibits NLRP3 inflammasome activation in human cells.** (A) Plasmids encoding VgrG2b C-terminus and the indicated human proteins were co-transfected into HEK293T cells for 24 h, followed by immunoblotting with antibodies against the indicated proteins. (B) Human macrophage cells were infected with lentiviruses encoding a control vector or VgrG2b C-terminus and primed with 1 μg/ml LPS for 3 h, followed by stimulation of 10 μM nigericin for 30 min, 0.5 μM gramicidin for 1 h and 2 mM ATP for 30 min. Cells were lysed and immunoblotted as indicated. (C–F) Wild-type BMDM cells were primed with 1 μg/ml LPS for 3 h, followed by stimulation of 10 μM nigericin with or without the transfection of VgrG2b C-terminus proteins using cell-penetrating peptides (CPP) for 30 min. Cells were lysed and immunoblotted (C). Cell culture supernatants were collected for an ELISA assay to determine the secreted IL-1β protein levels (D). Cytotoxicity was determined by LDH release assay in cell culture supernatants (E). Cell viability was determined by an ATP quantification assay in cell pellets (F). (G) FLAG-NLRP3 was purified from HEK293T cells using anti-FLAG-conjugated beads. NLRP3-containing beads were incubated with 100 μCi[γ-^32^P]ATP in the presence of 100 ng/ml VgrG2b C-terminus. Beads were washed with PBS and the associated radioactive ATP was determined by liquid scintillation. (H) Wild-type BMDM cells were primed with 1 μg/ml LPS for 3 h, followed by stimulation of 10 μM nigericin with or without the transfection of VgrG2b C-terminus proteins using cell-penetrating peptides (CPP) for 30 min. Cells were lysed and immunoprecipitated with antibody against NLRP3. The amount of mitochondrial DNA in the precipitates was examined through quantitative PCR. (I) Plasmids encoding VgrG2b C-terminus and NEK7 were co-transfected into HEK293T cells for 24 h, followed by immunoprecipitation with a control IgG or antibody against Myc. Precipitates were immunoblotted as indicated. (J) Plasmids encoding VgrG2b C-terminus, NEK7 and NLRP3 were co-transfected into HEK293T cells for 24 h, followed by immunoprecipitation with a control IgG or antibody against NLRP3. Precipitates were immunoblotted as indicated. (K) Recombinant VgrG2b C-terminus proteins were introduced into BMDM cells with the help of cell-penetrating peptides (CPP) for 6 h. Cells were then transfected with 4 μg/ml flagellin with the help of lipofectin for 4 h. Cells were lysed and immunoprecipitated with antibodies against the indicated proteins. Data were shown as means±SD. For S5D, S5E and S5F, data of three independent experiments were calculated. Experiments were repeated three times with similar results.

**Supplementary Figure 6. Activation of NLRP3 inflammasome ameliorates *P. aeruginosa* infection.** (A) Scheme for design of a VgrG2b C-terminus peptide. VgrG2b C-terminus was added with an N-terminal TAT sequence, followed by myristoylation. (B–E) BMDM cells were primed overnight with 1 μg/ml Pam3CSK4, followed by transfection of 2 μg/ml LPS using DOTAP with or without the incubation of ΔRetS PAO1 at an MOI of 30 and 2 μg/ml TAT-peptide for 2 h. Cells were then supplemented with fresh medium containing 100 μg/ml Gentamycin. Cells were lysed and immunoblotted as indicated 16 h post infection (B). Cell culture supernatants were collected for an ELISA assay to determine the secreted IL-1β protein levels 16 h post infection (C). Cytotoxicity was determined by LDH release assay in cell culture supernatants 16 h post infection (D). Cell viability was determined by an ATP quantification assay in cell pellets 16 h post infection (E). (F–I) Human macrophages were primed overnight with 1 μg/ml Pam3CSK4, followed by transfection of 2 μg/ml LPS using DOTAP with or without the incubation of ΔRetS PAO1 at an MOI of 30 and 2 μg/ml TAT-peptide for 2 h. Cells were then supplemented with fresh medium containing 100 μg/ml Gentamycin. Cells were lysed and immunoblotted as indicated 16 h post infection (F). Cell culture supernatants were collected for an ELISA assay to determine the secreted IL-1β protein levels 16 h post infection (G). Cytotoxicity was determined by LDH release assay in cell culture supernatants 16 h post infection (H). Cell viability was determined by an ATP quantification assay in cell pellets 16 h post infection (I). Data were shown as means±SD. For S6C, S6D, S6E, S6G, S6H and S6I, data of three independent experiments were calculated. Experiments were repeated three times with similar results.
