## Supplementary figures and images for "A VgrG2b fragment cleaved by caspase-11/4 promotes *Pseudomonas aeruginosa* infection through suppressing the NLRP3 inflammasome"

### Support Figure 1

#### Supplementary Figure 1

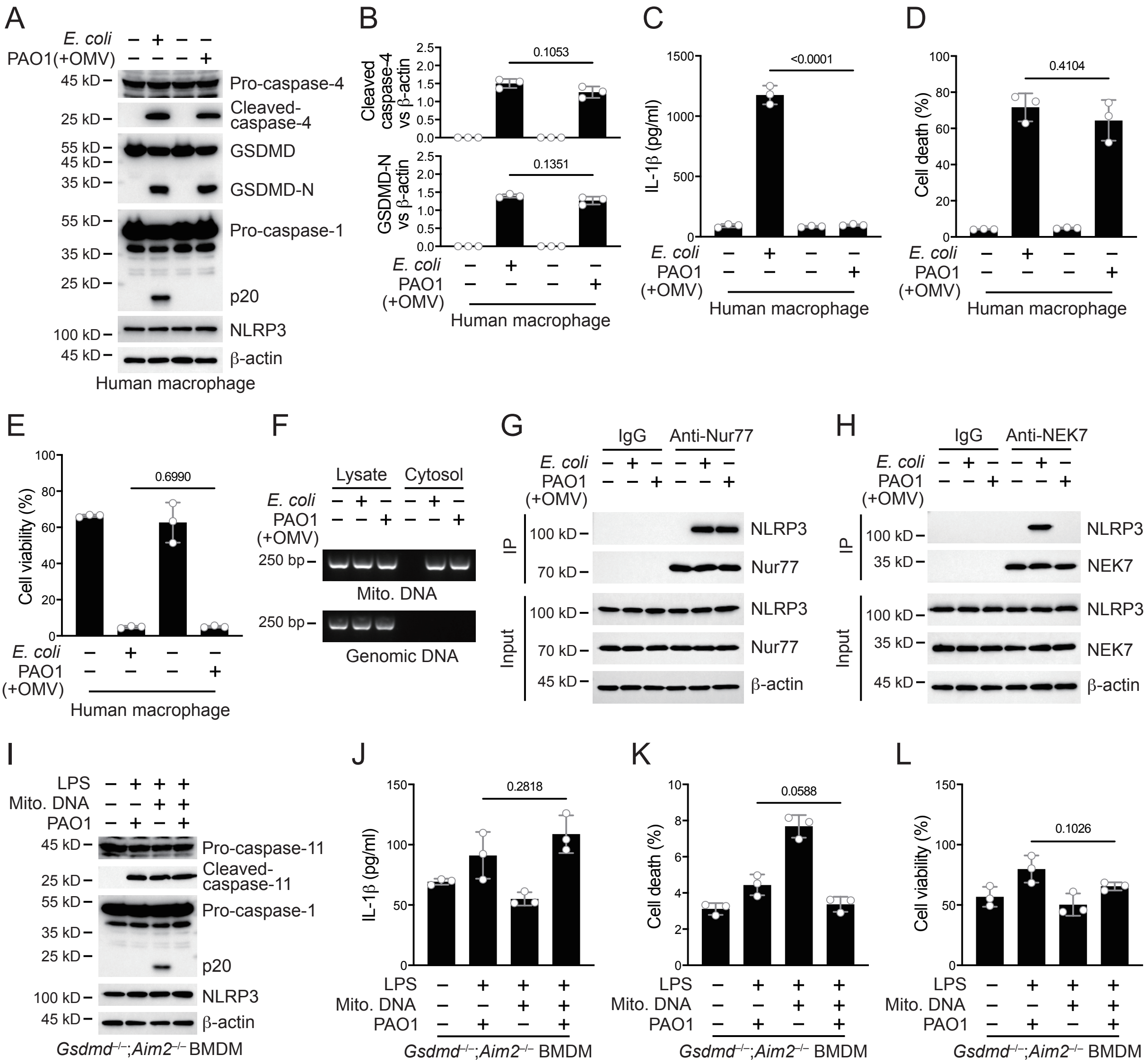

### Support Figure 2

#### Supplementary Figure 2

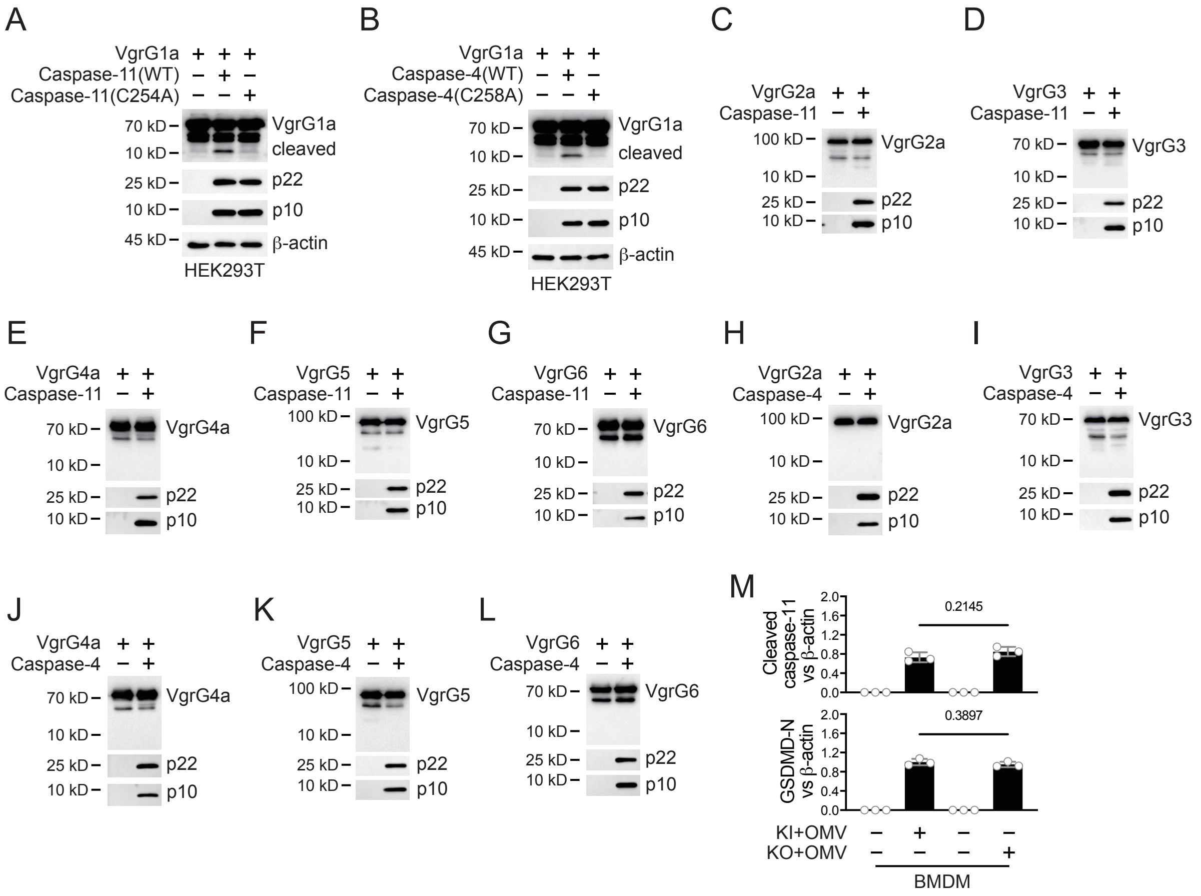

### Support Figure 3

#### Supplementary Figure 3

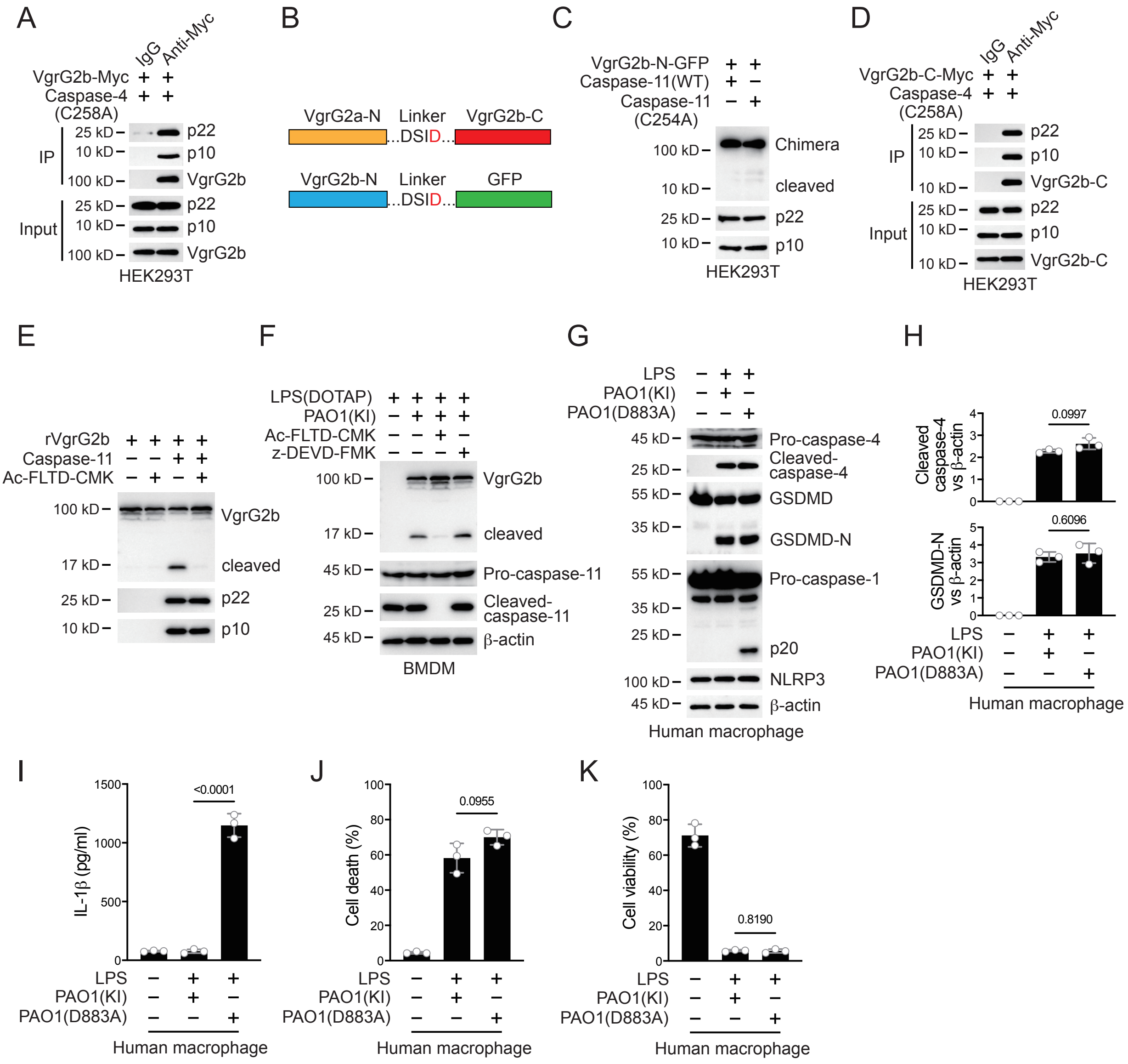

### Support Figure 4

#### Supplementary Figure 4

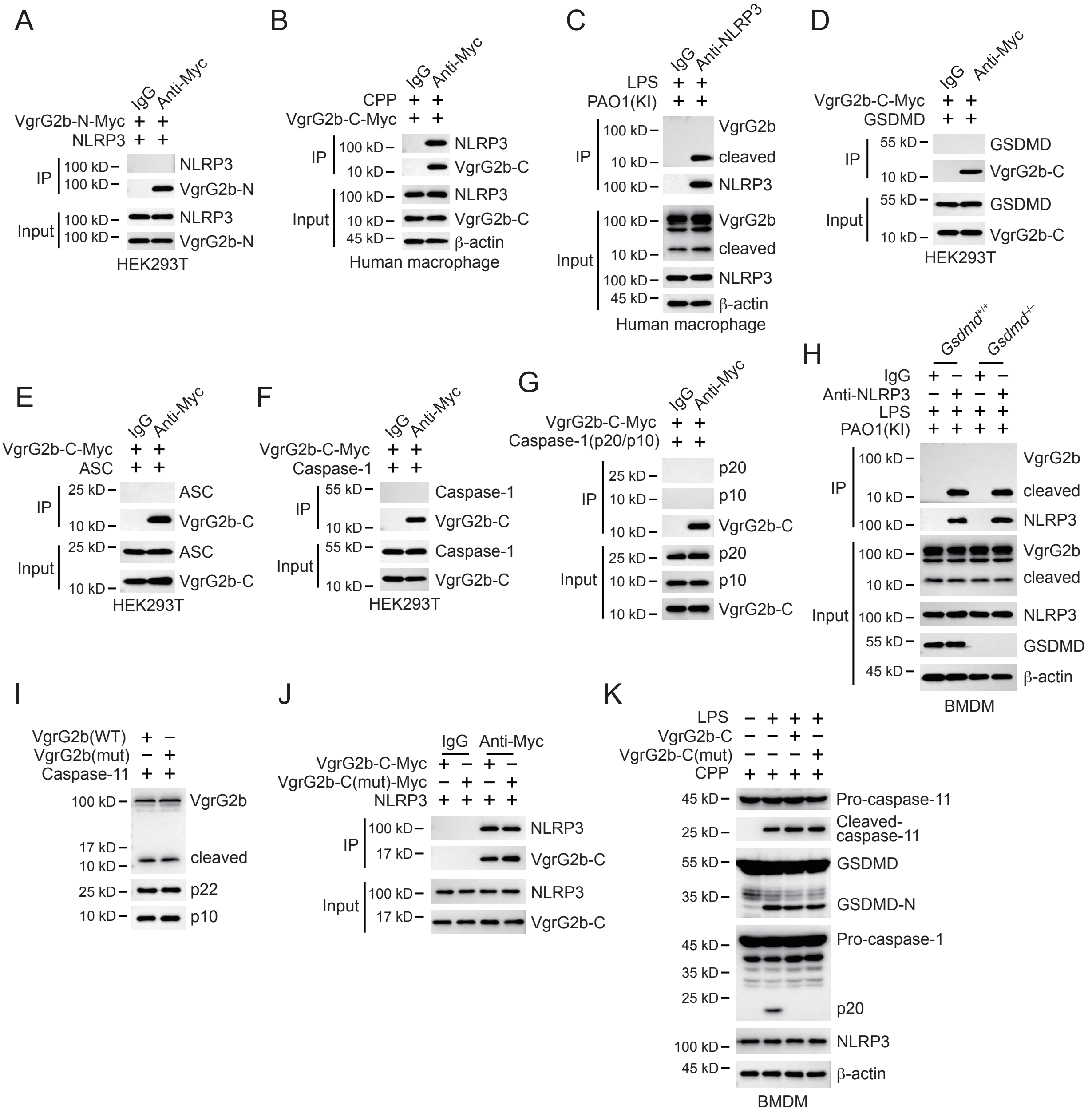

### Support Figure 5

#### Supplementary Figure 5

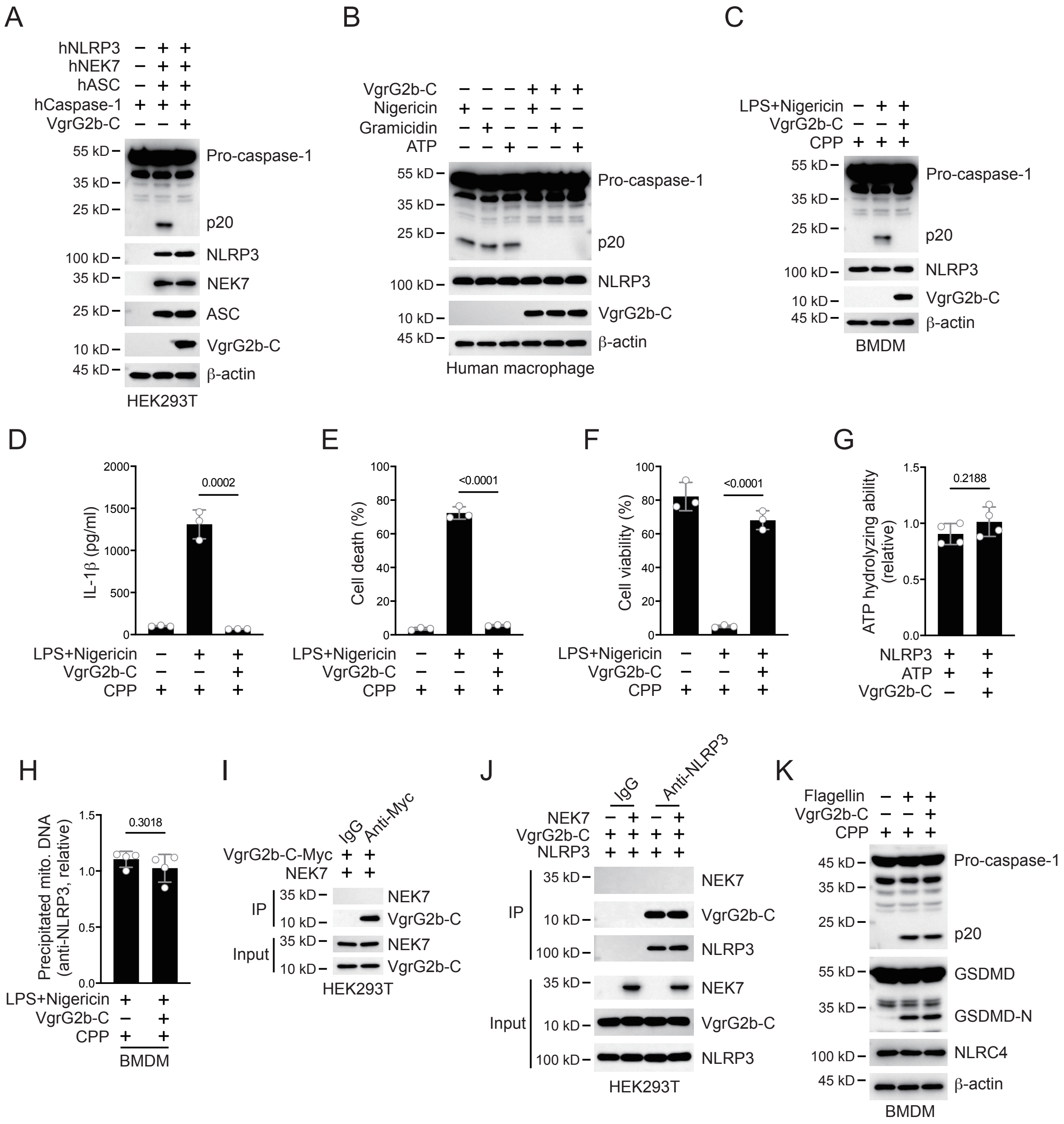

### Support Figure 6

#### Supplementary Figure 6

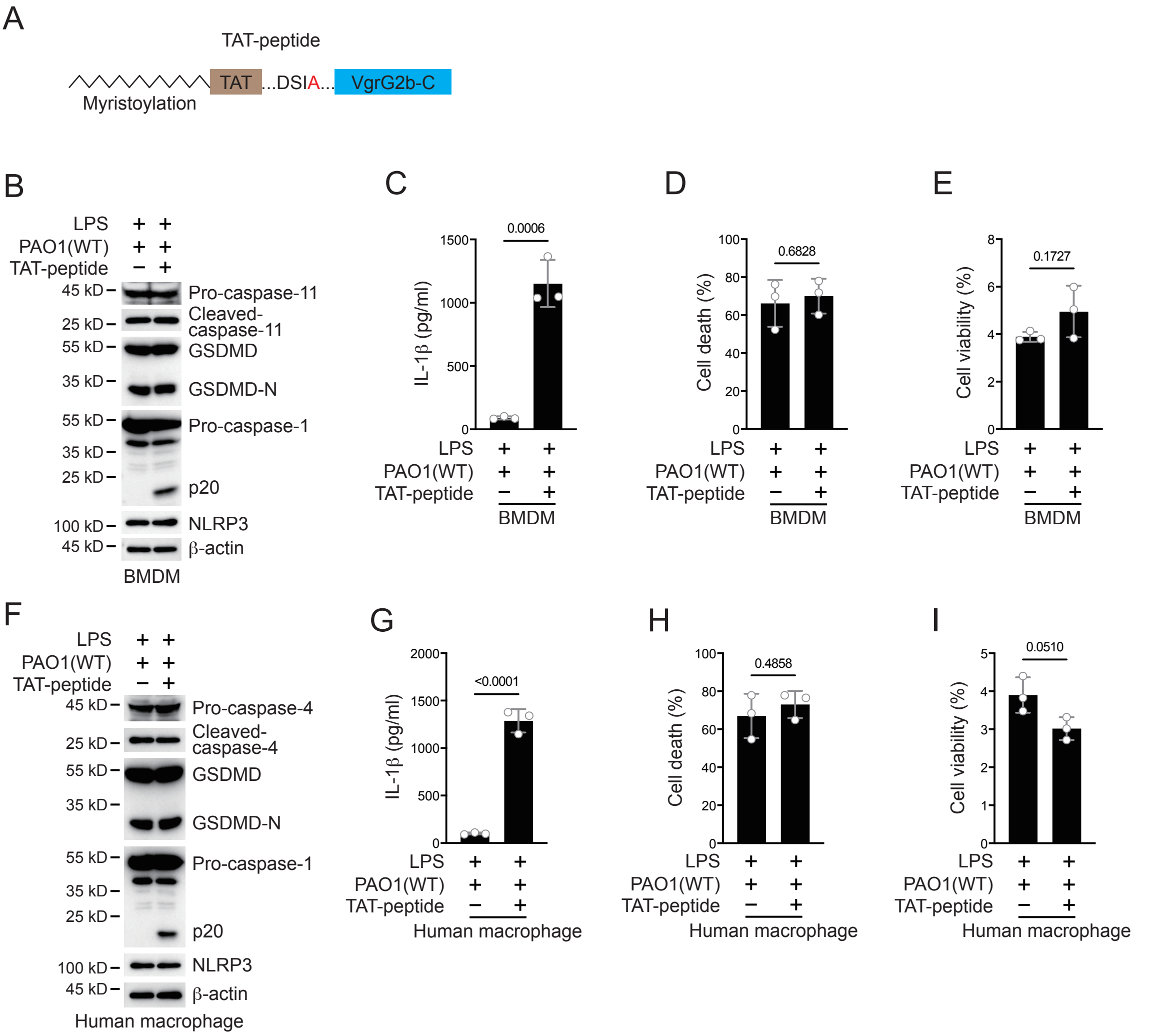
